# Parametric modulation of a shared midbrain circuit drives distinct vocal modes in a singing mouse

**DOI:** 10.1101/2025.04.04.647309

**Authors:** Xiaoyue Mike Zheng, Clifford E. Harpole, Martin B. Davis, Arkarup Banerjee

## Abstract

The ability of neural circuits to generate multiple outputs is critical for behavioral flexibility. Here, we leverage the rich vocal behavior of the singing mouse (*Scotinomys teguina*) to investigate the organizational logic of multifunctional motor circuits. We show that two distinct vocal modes—soft, unstructured ultrasonic vocalizations (USVs) for short-range and loud, rhythmic songs for long-range communication—arise not via parallel pathways but through shared brainstem phonatory circuitry involving the caudolateral periaqueductual gray (clPAG). Using a three-parameter linear model of song rhythm, we demonstrate that synaptic silencing of clPAG progressively alters song duration through a single parameter controlling its termination. This parameter also explains sexual dimorphism in songs, identifying clPAG as a key locus for driving natural behavioral variability. Our findings reveal how parametric modulation of a central circuit node can produce distinct behavioral modes, providing a mechanistic basis for rapid behavioral evolution in mammals.

## Introduction

Behavioral flexibility—the ability of animals to produce different actions depending upon context—is essential for survival. How the brain generates this flexibility remains a fundamental question in neuroscience. Distinct behavioral modes can be produced by dedicated motor circuits (1–3) or instantiated through shared circuits capable of operating at different functional regimes (4–7). Determining such organizational logic is critical for understanding not only the function but also the evolution of neural circuits. For instance, repurposing of shared, multifunctional circuits may drive rapid behavioral evolution, but the specific neural mechanisms underlying this process remain obscure, especially in mammalian brains.

Vocal communication, which requires coordinated respiratory, laryngeal, and orofacial motor control in different behavioral contexts, provides an ideal framework to investigate the rules of multifunctional circuit function (8–12). Singing mice (*Scotinomys teguina*), known for their distinctive songs used for turn-taking, are an attractive model system to explore this issue (13–16) (**Video S1**). We have previously shown that this behavior is dependent upon motor cortical function (17, 18). However, the downstream subcortical mechanisms that drive vocalizations remain unexplored. The periaqueductal gray (PAG), which projects to brainstem phonatory and respiratory networks, has been established as a necessary node for vocal production across vertebrate evolution (11, 19, 20). As we will show, the rich vocal behaviors of the singing mice and their control by the midbrain PAG provide an ideal testbed for addressing whether distinct vocal behaviors are generated by separate dedicated pathways or parametric modulation of a common circuit.

In this study, we address three specific questions: Do singing mice employ distinct vocal modes across different social contexts? If so, what are the acoustic characteristics and usage patterns of these modes? Do these distinct vocal behaviors emerge from separate neural pathways or from shared circuits operating in different regimes? While songs have been well studied, it remains unclear whether they represent their only major form of vocal communication. Additional vocalizations have been observed during social encounters (15, personal communication with Steven Phelps), yet their acoustic features, prevalence, and neural mechanisms remain unknown. By combining a novel behavioral paradigm with acoustic analysis, mathematical modeling, peripheral sound production experiments, and targeted manipulations of the caudolateral periaqueductal gray (clPAG), we report that singing mice use shared neural circuits for phonation operating under different amplitude modulation (AM) and frequency modulation (FM) regimes to generate categorically distinct vocal behaviors.

## Results

### Discovery of distinct vocal modes using PAIRId

An accurate and quantitative description of behavior is essential for investigating neural circuit mechanisms. We begin by measuring the vocal repertoire of individual singing mice during close-range social interactions. One major challenge in bioacoustics has been the assignment of quiet vocalizations produced in close proximity to one another. This has led to the development of methods such as wearable microphones or microphone arrays with subsequent sound localization using neural networks (21–25). Here, we took a complementary approach and developed a novel behavioral paradigm—PAIRId (**P**artial **A**coustic **I**solation **R**eveals **Id**entity)—where two animals placed in separate acoustically dampened enclosures equipped with private microphones and cameras can interact across a perforated plane (**Figure 1A, Figure S1A, Video S2**).

**Fig. 1.**
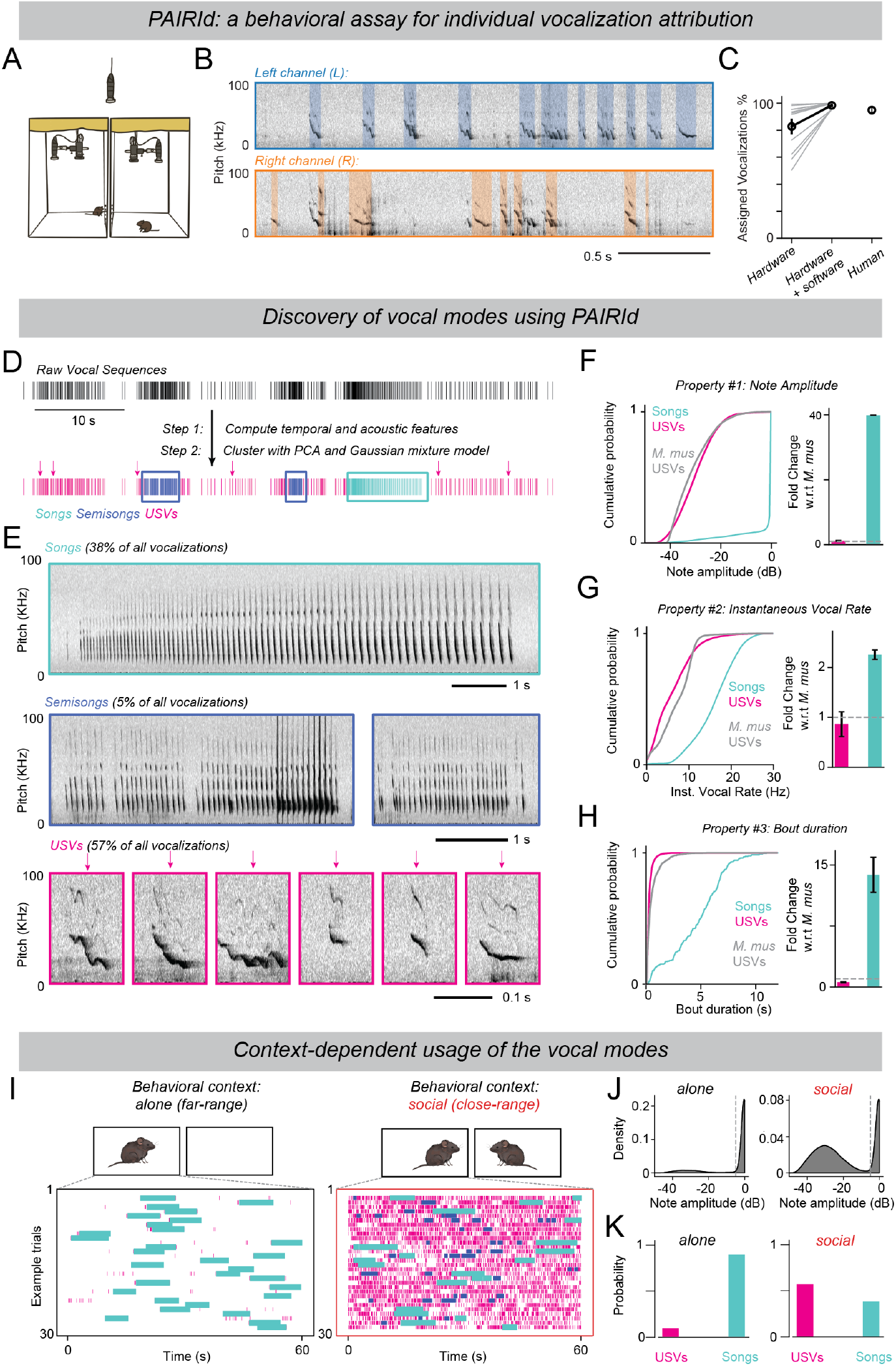
Discovery of distinct vocal modes of singing mice using PAIRId. **(A)** Schematic of the PAIRId behavioral paradigm, where two mice housed in separate acoustically dampened enclosures—each equipped with its own microphone and camera—interact across a perforated plane. **(B)** Example audio data from two singing mice interacting in PAIRId, where blue and orange boxes denote vocalizations detected and attributed to individual mice. **(C)** Proportion of vocalizations attributed to individual mice using hardware acoustic isolation (82.8 ± 5.0%) and the full analysis pipeline (98.4 ± 0.6%, n = 12 sessions of 5 hours each), compared with human annotations (94.9 ± 1.7%, n = 4 hours in separate sessions). **(D)** Schematic for identifying vocal modes from temporal sequence of vocalizations. *Top*: Raster plot illustrating vocalizations of an animal during an example 60-second period, with each line marking the onset of a vocalization. *Bottom*: The same raster plot, with lines color-coded by the vocal modes (songs, cyan; semisongs, blue; USVs, magenta) identified by combining temporal stereotypy and loudness features. **(E)** Example spectrograms for each of the three identified vocal modes—songs, semisongs, and USVs—highlighted in (D). To account for the large loudness difference among vocalizations, the song spectrogram was computed from low-gain audio recording whereas the semisong and USV spectrograms were computed from high-gain audio recording. **(F)** Loudness of notes in different vocal modes. *Left*: Cumulative distribution of note loudness for each vocal mode (singing mouse USV: n = 26,246 notes from 10 mice, -29.78 ± 0.05 dB; singing mouse song: 17,681 notes from 10 mice, -2.27 ± 0.05 dB; laboratory mouse USV: n = 7,295 notes from 3 mice, -30.82 ± 0.09 dB). *Right*: Fold change of median note loudness per mouse relative to the median of laboratory mouse USVs (singing mouse USV: 1.1 ± 0.2, P > 0.05; singing mouse song: 39.9 ± 0.0, P = 0.007). **(G)** Same as (F), but for instantaneous note rate (singing mouse USV: 6.26 ± 0.03 Hz, fold change 0.9 ± 0.2, P > 0.05; singing mouse song: 16.16 ± 0.04 Hz, fold change 2.3 ± 0.1, P = 0.007; laboratory mouse USV: 7.24 ± 0.05 Hz). **(H)** Same as (F), but for vocal bout duration (singing mouse USV: 0.31 ± 0.00 s, fold change 0.6 ± 0.1, P = 0.007; singing mouse song: 4.64 ± 0.15 s, fold change 13.8 ± 2.2, P = 0.007; laboratory mouse USV: 0.56 ± 0.02 s). **(I)** Raster plots of 30 segments with the highest vocal activity when the mouse is alone (*left*) vs. during close-range social interactions in PAIRId (*right*). Each row represents one minute, and each tick indicates a vocalization, color-coded by its assigned mode. **(J)** Distribution of loudness of all vocalizations when mice are alone (*left*, n = 23,028 notes) versus when they interact socially in PAIRId (*right*, n = 46,027 notes). **(K)** Proportion of songs and USVs across these two behavioral contexts (same as in (J)). Unless stated otherwise, values reported are Mean ± SEM and hypothesis testing was performed using Mann-Whitney U test.

To identify vocalizations of individual animals, we leverage both hardware-level acoustic isolation and software-level signal processing (**Figure 1B**). Briefly, vocalizations detected exclusively in one channel due to acoustic dampening are assigned directly to that animal (82.8 ± 5.0%). For the remaining detections with temporal overlap across channels, we compare spectrotemporal properties to distinguish coincident vocalizations and acoustic bleed-through (**Figure 1B, Figure S1B, Methods**). This approach allows for the attribution of almost all vocalizations (98.4 ± 0.6%, 12 male-female pairs) with high accuracy (F1 score = 0.87 ± 0.02, 4 hours), comparable to human annotations (**Figure 1C, Figure S1C**). Taken together, PAIRId allows us to quantify the vocalizations of individual singing mice during social encounters, providing the necessary foundation for exploring underlying neural mechanisms.

Using PAIRId, we found that the singing mouse vocal repertoire is much richer than previously anticipated. Both male and female singing mice interact robustly within a few body lengths of each other, producing numerous vocalizations in correlated bouts (**Figure S1D–G**). It further revealed the complexity of temporally sequenced vocal streams produced by individual singing mice (**Figure 1D**). To quantitatively describe their vocal repertoire, we analyzed the temporal rhythm of vocal bouts (**Figure S2A, Methods**). We identified rhythmic patterns—where mice produce notes at steady or smoothly varying rates—by first detecting short, contiguous sequences with high temporal stereotypy. These vocal segments were then hierarchically merged into longer sequences, which we clustered using a Gaussian mixture model to reveal distinct rhythms (**Figure S2B–E**). In addition to these rhythmic vocal sequences, we also observed substantial differences in loudness of individual notes (**Figure S2G *bottom***). Combining temporal characteristics and loudness information, we define two major vocal modes—songs and ultrasonic vocalizations (USVs) (**Figure 1E, Figure S2F– H**). We also observed a rarer vocal mode (semisongs, 5% of vocalizations) with acoustic properties intermediate between USVs and songs (**Figure S2I**). Given its rarity, we subsequently focused on songs and USVs, which together comprise 95% of all vocalizations.

To place the two vocal modes of singing mouse in a phylogenetic context, we compare them with vocalizations of other rodents. Laboratory mouse (*Mus musculus*) USVs serve as an ideal comparative reference due to their thorough characterization in the literature (26–28). We found that singing mouse USVs have overlapping acoustic properties with laboratory mouse USVs—including lower amplitude, lower temporal stereotypy, shorter bouts (**Figure 1F–H *magenta***), and variable note shapes (**Figure 1E *magenta***). In contrast, songs are much louder, have faster tempos, and contain many more stereotyped notes over much longer durations (**Figure 1F–H *cyan***). We conclude that songs have acoustic characteristics in both amplitude and temporal domains that are categorically distinct from the USVs of both species.

This acoustic distinction is reflected in context-dependent usage. When alone, singing mice produce songs almost exclusively (**Figure 1I–K, Figure S2B**). During close-range interactions, the vocal repertoire becomes more complex, composed of both songs and quieter USVs (**Figure 1I *right*, Figure S2C**). The overall vocal loudness decreases considerably (**Figure 1J**), with a substantial increase in the proportion of USVs (**Figure 1K**). Notably, we found that singing mice can switch between these vocal modes in rapid succession (**Figure S3**). We conclude that singing mice employ two dominant vocal modes for different social contexts. Softer USVs, which attenuate sharply over distance, are used predominantly during close-range social interactions, consistent with their role in short-range communication across a variety of rodent species (29). In contrast, songs are produced when mice are alone or engaged in vocal turn-taking with conspecifics under visual occlusion, as we have previously shown (18), supporting their function in long-range communication.

### A linear model of the song rhythm

Singing mouse songs are strikingly divergent from the USVs in their temporal stereotypy. To capture this stereotyped rhythmic structure, we sought to develop a mathematical model describing the temporal progression of individual notes using a compact set of parameters. Songs are composed of a series of increasingly longer notes produced over several seconds (**Figure 2A**). We serendipitously observed that during song progression, instantaneous note rate *r*(*i*)—defined as the inverse of inter-note interval—varies linearly with note index *i*. A song always begins with notes emitted at a high rate, which steadily decreases throughout the song, finally ending at a much lower rate (**Figure 2B**). A linear model with just three parameters (start rate *r*_max_, slope *m*, and stop rate *r*_min_) adequately describes the instantaneous note rate throughout the song:

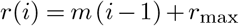

for *i* = 1, 2,…, *N −* 1 where *N* is the number of notes in a song (**Figure 2B, Methods**). This simple generative model reliably captures the temporal progression of songs across our entire dataset (**Figure 2C**). Moreover, song duration *T* estimated by integrating the instantaneous rate in the time domain matches the measured song durations accurately across all animals (**Figure 2D**):

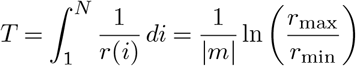

**Fig. 2.**
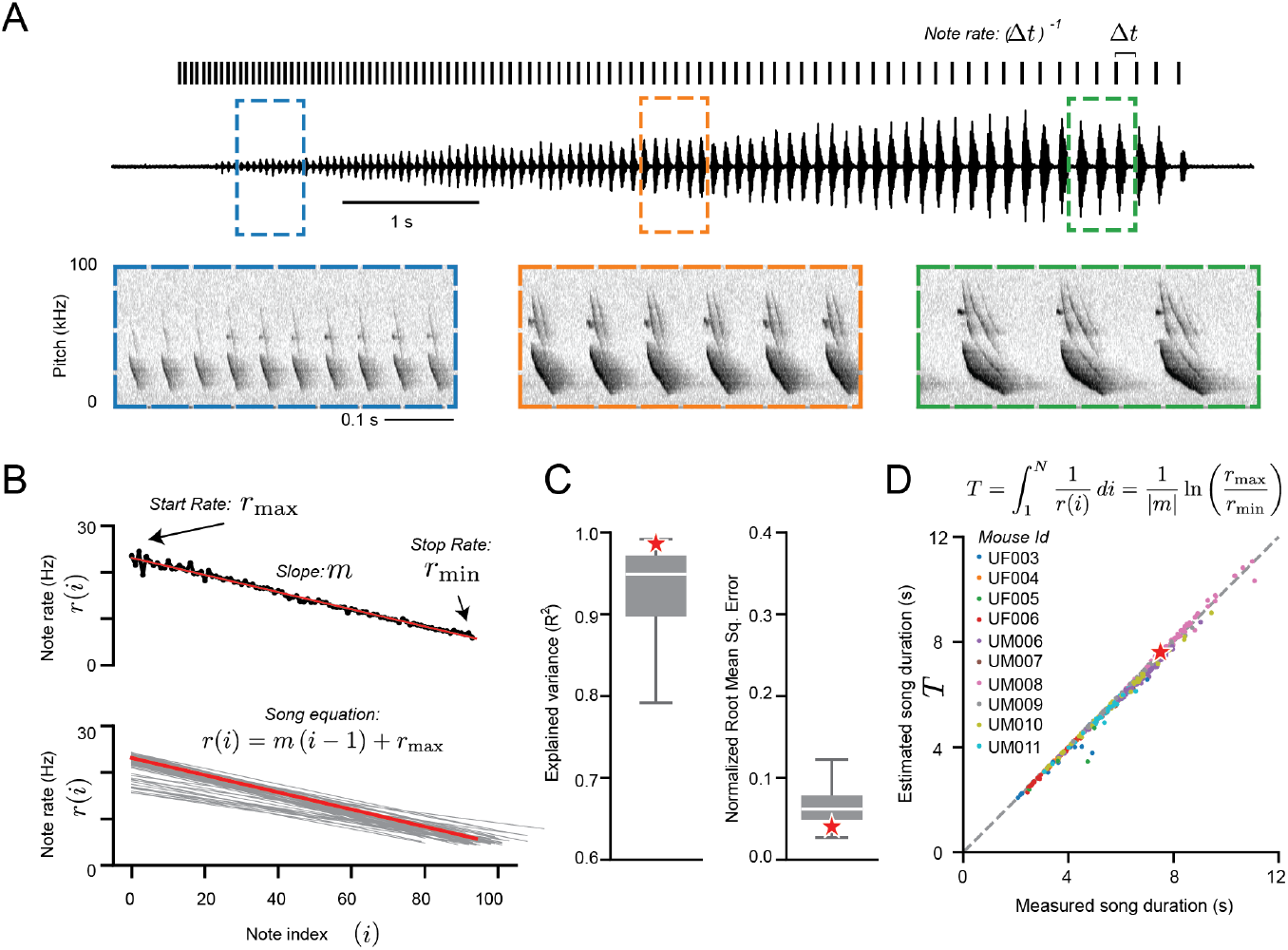
A linear model of the song rhythm. **(A)** An example singing mouse song. *Top*: Raster plot of the song, with each line marking the onset of a song note. The annotation indicates that the instantaneous note rate is calculated as the inverse of the interval between note onsets. *Middle*: Audio waveform of the song. *Bottom*: spectrograms of the highlighted snippets of the song. **(B)** Rhythm trajectories of example songs. *Top*: The trajectory of the example song from (A) is plotted with each note’s instantaneous rate against its sequential index; the red line shows the linear model fit. *Bottom*: Model fits of rhythm trajectories for all 60 songs produced by a single mouse, with the red line indicating the example song. **(C)** Performance of the linear model fit (n = 456 songs from 10 mice across alone and social contexts, *R*^2^ = 0.90 ± 0.01, normalized root mean squared error (NRMSE) = 0.07 ± 0.00). Star represents the example song from (A). **(D)** Model-predicted versus measured song duration (n = 456 songs from 10 mice).

Thus, this linear model captures the stereotyped rhythm of singing mouse songs and explains the observed variability through just three patterning parameters.

### Peripheral mechanisms for distinct vocal modes

As we have shown above, singing mouse songs are rhythmic and loud, but their USVs are much softer with less temporal stereotypy. What might be the neural mechanisms driving these two distinct vocal modes? Given their categorical differences, one possibility is that these two vocal modes are largely driven by parallel motor pathways. Alternatively, the presence of the intermediate semisongs and the smooth transitions between songs and USVs might imply that the two distinct vocal modes share a common motor pathway operating in two different regimes (**Figure S2I, Figure S3**). Building on the current understanding of the mammalian vocal motor hierarchy (11, 12), we tested these alternative models at three different levels: peripheral mode of sound production, phonation-respiration coupling, and mechanisms of vocal gating by the PAG in the midbrain.

We first determined whether the biophysical mechanism for sound production differs between songs and USVs (**Figure 3A–G**). Rodents produce sounds using their larynx in two ways: by vocal fold vibrations (e.g. squeaks in laboratory mouse) or by generating aerodynamic whistles within the larynx (e.g. USVs in laboratory mouse) (30). While the precise details of the laryngeal and aerodynamic mechanisms remain under investigation (31–33), the two broad categories can be distinguished by simply changing the density of the surrounding air (30, 34, 35). If sounds are produced by a whistle mechanism, the fundamental frequency (F0) will depend on air density and increase in a less dense medium. In contrast, if sounds are produced by a vibrational mechanism, F0 is expected to remain unchanged. To test these alternatives, we replaced air in the behavioral enclosure with a helium-oxygen mixture (heliox; 80% helium, 20% oxygen). Under heliox, F0 of both USVs and song notes increased significantly— and by similar ratios—consistent with an aerodynamic whistle mechanism (**Figure 3B–E**). Crucially, we verified that singing mice were also able to vocalize via a vibrational mechanism, as the F0 of their squeaks recorded concurrently remained stable (**Figure 3F–G**). These findings demonstrate that both USVs and songs are produced by a similar whistle mechanism at the vocal periphery.

**Fig. 3.**
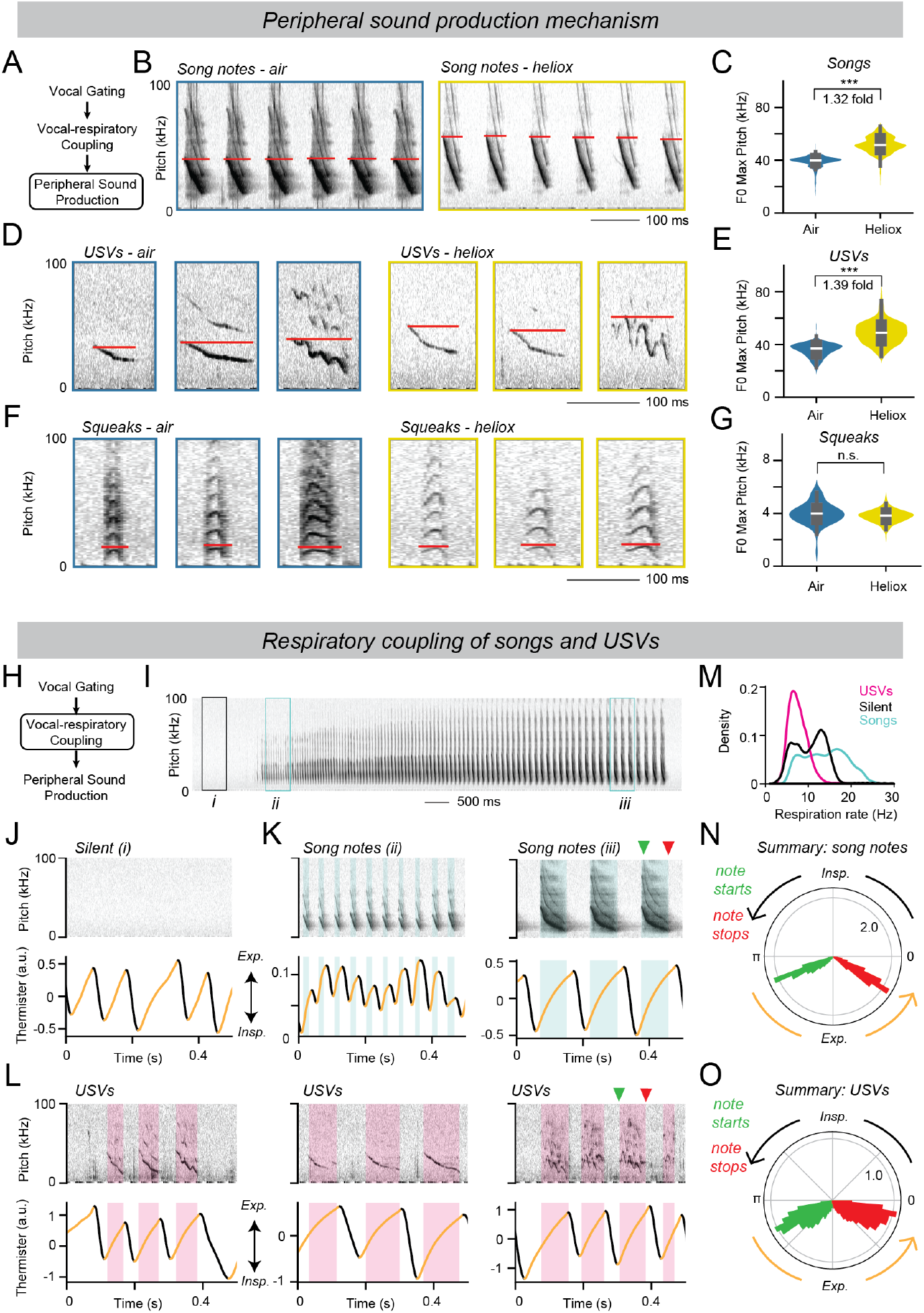
Shared peripheral vocal production mechanisms between songs and USVs. **(A**) Schematic of the vocal motor hierarchy, highlighting the level addressed by the laryngeal phonation mechanism experiment. **(B**) Example spectrograms of song notes produced in air and in helium-oxygen mixture (heliox), with the red lines illustrating the maximum of the fundamental frequency (F0). **(C)** The maximum F0 of song notes is significantly higher in heliox (51.7 ± 0.1 kHz, n = 1,648 notes) than in air (39.0 ± 0.1 kHz, n = 2,262 notes; P = 0.0), consistent with a whistle mechanism. **(D** and **E)** Same as (B) and (C), but for USVs instead of song notes, showing an increase in F0 in heliox (35.8 ± 0.2 kHz, n = 400 notes in air; 49.1 ± 0.4 kHz, n = 400 notes in heliox; P = 1.38e-94), also consistent with a whistle mechanism. **(F** and **G)** Same as (B) and (C), but for squeaks instead of song notes, showing F0 in heliox is not significantly higher than in air (4.0 ± 0.1 kHz, n = 75 notes in air; 3.8 ± 0.0 kHz, n = 208 notes in heliox; P > 0.05), consistent with a vibration mechanism. **(H**) Schematic of the vocal motor hierarchy, highlighting the level addressed by the vocal-respiratory coordination experiment. **(I)** Spectrogram of a song preceded by a silent period, recorded with simultaneous thermistor-based respiration monitoring. **(J)** Example respiration dynamics during silence (box *i* in (I)), with black indicating inhalation and orange indicating exhalation. **(K)** Same as (J), but during song production (boxes *ii* and *iii* in (I)). Song notes, shown in cyan, occur during exhalation. **(L)** Same as (J), but for when the mouse makes USVs. USVs, shown in magenta, also occur during the exhalation phase. **(M)** Distribution of respiration rate when the mouse is silent (black), making songs (cyan), or making USVs (magenta). **(N)** Polar plots showing distribution of note onsets (green) and offsets (red) relative to the respiratory cycle (inspiration: 0–π; exhalation: π–2π) for song notes (onset 3.71 ± 0.01 rad, offset 5.74 ± 0.00 rad, n = 2,384 notes). **(O)** Same as (N), but for USVs (onset 3.93 ± 0.01 rad, offset 5.70 ± 0.01 rad, n = 1,910 notes).

Moving up the vocal motor hierarchy, we next examined the coupling between phonation and respiration during songs and USVs (**Figure 3H–O**). We implanted a thermistor in the nasal cavity to continuously monitor temperature changes as a proxy for respiration **(Figure 3I–L)**. We found each song note was associated with a single respiration cycle, with phonations occurring exclusively during exhalation (**Figure 3K, N**), consistent with previous findings using a different method (18). Therefore, the temporal patterning of song notes arises from the rhythmic structure of respiration it-self. Similar to song notes, each USV, regardless of duration, is produced during exhalation with a similar phase relationship (**Figure 3L, O**). This is in line with previous literature demonstrating that laboratory mouse USVs are tightly coupled to underlying respiratory cycles and are produced exclusively during exhalations (26, 36). While the respiration rates during USVs fall within the range observed during silent, non-vocal periods, respiratory rates needed for singing are substantially higher and can easily exceed 20 cycles per second (**Figure 3M**). Our results demonstrate that the two strikingly distinct vocal modes—songs and USVs—share peripheral sound production mechanism as well as vocal-respiratory coupling. This suggests that in the singing mouse, the newly-evolved song mode has co-opted the phonatory mechanism of USV production, while operating it at substantially higher amplitude and frequency regimes for song motor control.

### Vocal gating by the caudolateral PAG

Given that USVs in singing mice and laboratory mice share similar acoustic properties, behavioral contexts, and peripheral sound production mechanisms, we hypothesized that their vocal gating mechanisms might also be conserved. The midbrain periaqueductal gray (PAG) has been established as a critical locus for innate behaviors, including vocalizations, across vertebrates (20). Specifically, neurons in the caudolateral portion of the PAG (clPAG) project to hindbrain phonation circuits (e.g., the nucleus retroambiguus (RAm), the intermediate reticular oscillator (iRO), and the pre-Bötzinger complex (preBötC), and are necessary and sufficient for USV production in laboratory mice (37–42). Therefore, we decided to examine the role of clPAG for vocal gating in the singing mouse.

We relied on gross morphological similarities to identify the homologous PAG subregion. Optogenetic activation of channelrhodopsin-2 (ChR2)-expressing clPAG neurons was sufficient to reliably elicit vocalizations in the singing mouse (**Figure 4A–B, Video S3**). Vocalizations began immediately after light onset and scaled with the duration of photostimulation (**Figure 4B–D**, n = 4 mice). Acoustic analyses revealed that these vocalizations resembled USVs in both amplitude and instantaneous vocal rates (**Figure 4E**). Therefore, clPAG stimulation is sufficient to evoke USVs in singing mice. Conversely, synaptic silencing of clPAG neurons with Tetanus toxin light-chain (TeLC) caused a severe reduction in USVs across all animals tested (**Figure 5A–B**, n = 5 mice). We conclude that clPAG neurons are both necessary and sufficient to generate species-typical USVs in the singing mouse. We also found that synaptic silencing of clPAG also eliminated songs; in effect, the animals were rendered mute (**Figure 5C**). This observation rules out a separate, parallel pathway for song production independent of the clPAG. Instead, it points to a shared vocal motor control circuit for both USVs and songs. Based on similarities in acoustic properties, behavioral context, peripheral production mechanisms, and sufficiency of eliciting USVs from the clPAG, we posit that USVs in singing mice and laboratory mice are homologous behaviors. This is despite superficial differences in pitch (USVs in laboratory mice have a much higher pitch), which presumably reflects differences in airway and laryngeal morphology, such as the size of the ventral pouch (43, 44). In contrast, songs represent a drastically divergent vocal behavior unique to the singing mice lineage, characterized by two key innovations in motor control: significantly higher amplitude and highly stereotyped temporal patterning over long bouts.

**Fig. 4.**
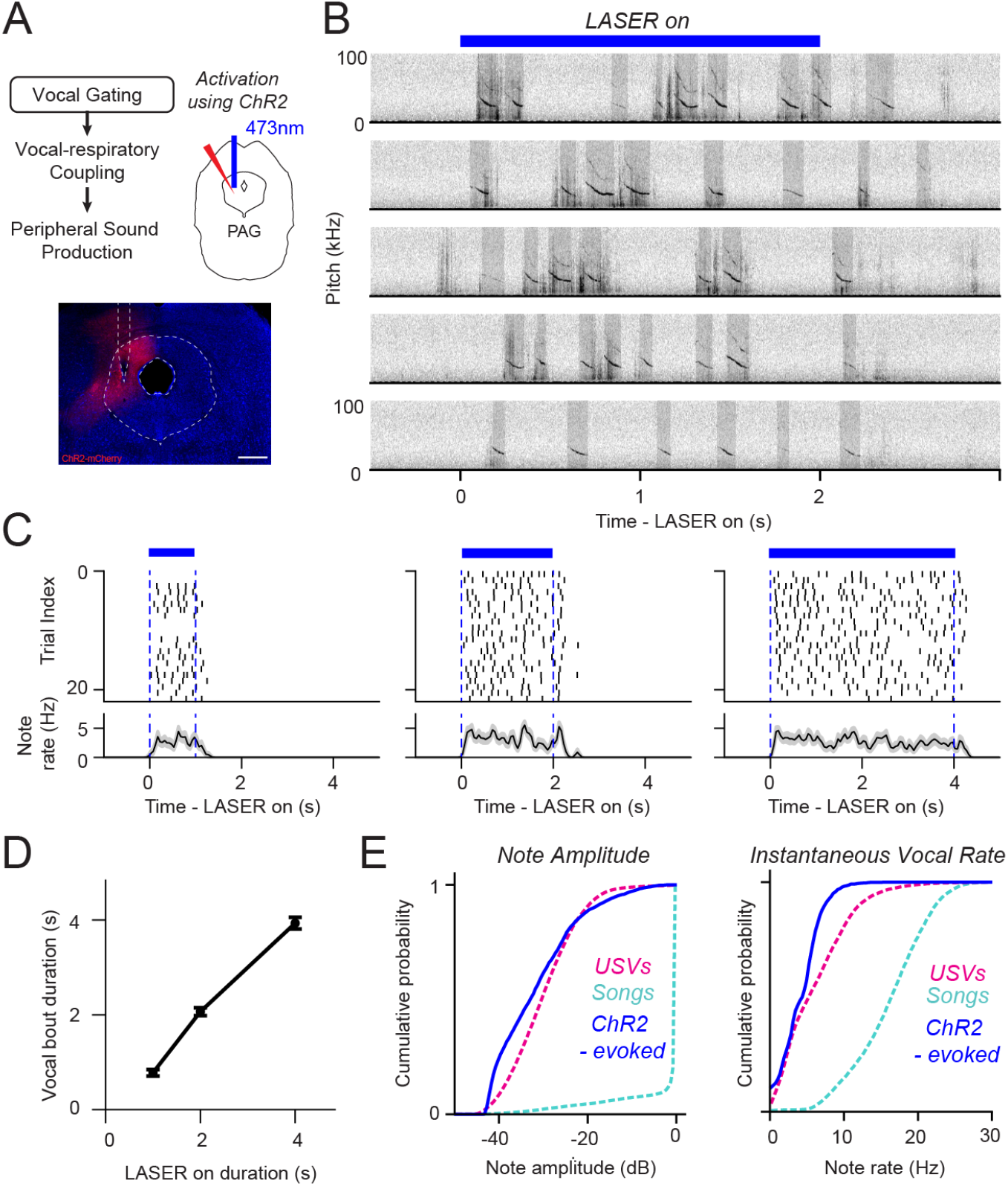
Activating midbrain clPAG is sufficient to elicit USVs in singing mice. **(A)** *Top left* : Schematic of the vocal motor hierarchy, highlighting the level addressed by the clPAG optogenetic activation experiment. *Top right* : Schematic of the experiment, showing the unilateral virus injection in the clPAG to express ChR2-mCherry, with the fiber implanted above. *Bottom*: Example image of the clPAG displaying virus expression and the fiber implant. **(B)** Spectrograms from five example trials of 2-second tonic optogenetic activation of the clPAG in an example singing mouse, showing evoked vocalizations during the stimulation period. **(C)** Vocalization raster and rate (Mean ± SEM) for optogenetic activations of 1 second (*left*), 2 seconds (*middle*), and 4 seconds (*right*) in an example mouse, demonstrating that vocalizations are elicited throughout the stimulation period in each condition. **(D)** Duration of the evoked vocalization bout compared with the duration of the optogenetic stimulation for all mice (1s stimulation: 0.78 ± 0.07 s, n = 86 trials; 2s stimulation: 2.07 ± 0.08 s, n = 85 trials; 4s stimulation: 3.93 ± 0.12 s, n = 85 trials; n = 4 mice). **(E)** Cumulative distributions of note amplitude (*left*) and instantaneous note rate (*right*) of optogenetically evoked vocalizations (amplitude: -31.4 ± 0.2 dB; rate: 4.27 ± 0.06 Hz; n = 2,063 notes) are similar to those of natural USVs (amplitude: -29.8 ± 0.0 dB; rate: 6.26 ± 0.03 Hz; n = 26,246 notes) and distinct from songs (amplitude: -2.3 ± 0.1 dB; rate: 16.16 ± 0.04 Hz; n = 17,681 notes).

**Fig. 5.**
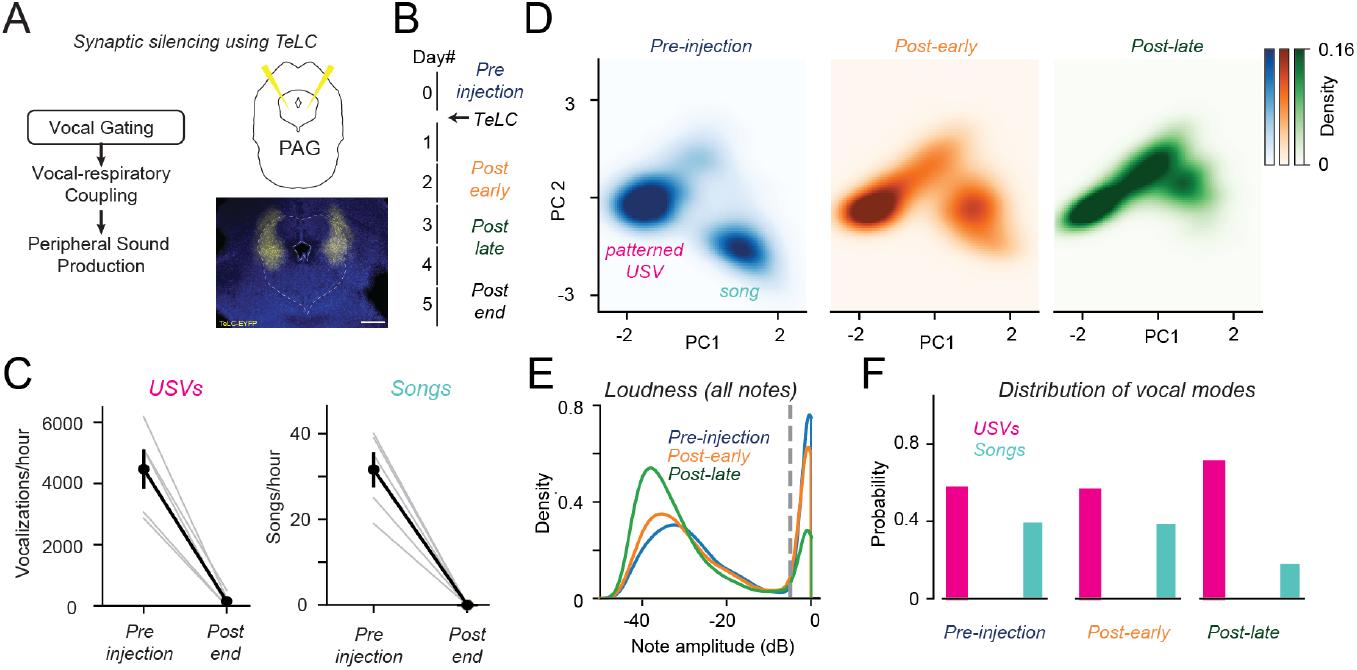
Synaptic silencing of the clPAG progressively degrades the vocal repertoire of singing mice. **(A)** *Left*: Schematic of the vocal motor hierarchy, highlighting the level addressed by the clPAG synaptic silencing experiment. *Top right* : Schematic of the experiment, showing the bilateral virus injection in the clPAG to express TeLC-EYFP. *Bottom right* : Example image of the clPAG displaying virus expression. **(B)** Experimental timeline of the clPAG silencing experiment with tetanus toxin light chain (TeLC). To summarize data across slightly different timelines across animals, the post-injection period for each animal was binned into two halves, with “post-early” covering the hours containing the first half and “post-late” the second half of all curated vocalizations. **(C)** Number of USVs (left; pre-injection, 4469 ± 645 notes/hour, post-injection: 157 ± 102 notes/hour, P = 0.008) and songs (right; pre-injection: 32 ± 4 songs, post-injection: 0 ± 0 songs, P = 0.008) before and after the virus injection (n = 5 mice), quantified as the number of vocalizations in a highly vocal hour before injection and 5-6 days after injection. **(D)** Distribution of vocal segments in the temporal feature space (see **Figure S2**) for curated hours before virus injection (*left*); 428 segments from 39,596 notes), during the post-early period (*middle*); n = 408 segments from 34,826 notes), and during post-late period (*right* ; n = 252 segments from 21,084 notes), showing a shift in density away from songs. **(E)** Distribution of loudness for all vocalizations before virus injection (n = 39,596 notes), during the post-early period (n = 34,826 notes), and during post-late period(n = 21,084 notes), showing an overall reduction in loudness. **(F)** Proportion of songs and USVs across these experimental timepoints (same as in (E)).

### Parametric control of song progression by the clPAG

How does the neural circuitry governing the elaborate songs differ from that of USVs? So far, we have shown that the mechanisms for phonatory control is shared between songs and USVs. However, given the drastic differences in loudness and rhythm during songs (compared to USVs), we hypothesized that song production involves driving the same circuit node (clPAG) in a higher amplitude and frequency regime. This would predict that progressive silencing of the clPAG would lead to graded disruptions in song quality. To test this possibility, we summarized data across all TeLC injected animals by binning each animal’s post-injection period into two equal halves: “post-early” (containing the first half of all analyzed vocalizations) and “post-late” (containing the second half). Consistent with our hypothesis, we observed a progressive decrease in both loudness and rhythmicity away from songs (**Figure 5D-E, Figure S4A**), leading to an increasing proportion of USVs (**Figure 5F**). Therefore, synaptic silencing of clPAG was associated with a progressive dialing down of both loudness and rhythmicity—the two acoustic features that distinguish the novel song mode compared to the ancestral USVs.

To better understand how clPAG influences song production, we used our analytical framework to examine the changes in song quality following TeLC-injection. Compared to pre-injection control songs, we observed a progressive reduction in note loudness in both example songs (**Figure 6A–B**) and across animals (**Figure 6C**). In the temporal domain, song duration also progressively decreased (**Figure 6A, D**). Using our linear model of song rhythm, we decomposed each song into its three underlying patterning parameters. We found that clPAG silencing led to a large increase in the stop rate (*r*_min_), while the other two parameters were only modestly affected (**Figure 6E, Figure S4B**). In fact, the relationship between song durations, *T*, and stop rates, *r*_min_, for all songs closely followed our model’s prediction: *T* would vary inversely with the negative logarithm of the *r*_min_ (**Figure 6F**). Thus, clPAG silencing shortened songs by causing earlier termination. In other words, male mice were no longer able to sustain their song motor pattern and append long, loud notes at the end of songs (**Figure 6G, Figure S4D**). As songs progressively deteriorated over days, they eventually broke into multiple, barely recognizable fragments before disappearing entirely—consistent with cumulative degradation of phonation, loudness, and rhythmic structure (**Figure S5**). Taken together with the necessity of the clPAG for producing both songs and USVs, we conclude that the song mode is produced not via a separate pathway but through amplitude modulation (AM) of individual notes and frequency modulation (FM) of motor patterning in the clPAG.

**Fig. 6.**
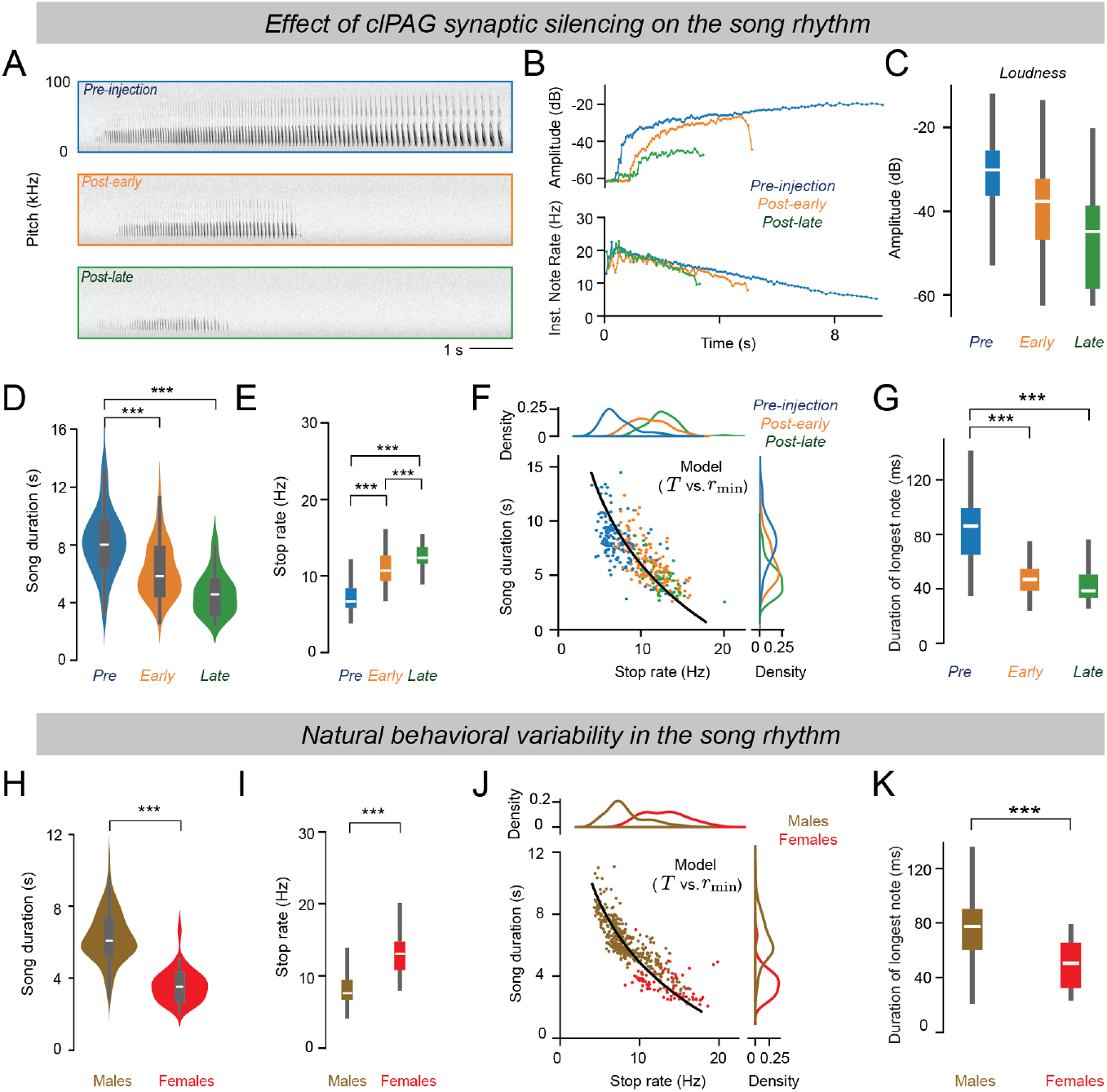
Parametric modulation of song patterning by the clPAG. **(A)** Spectrograms of example songs from one mouse before the PAG TeLC virus injection (*top*), during the post-early period (*middle*), and during post-late period (*bottom*). **(B)** Trajectories of note amplitude (*top*) and instantaneous note rate (*bottom*) for the example songs shown in (A). **(C)** Distribution of note loudness in low-gain audio for all song notes across different PAG TeLC experimental timepoints (n = 5 mice; pre-injection: n = 15,583 notes, -32.53 ± 0.09 dB; post-early: n = 13,490 notes, -40.60 ± 0.10 dB; post-late: n = 3,774 notes, -46.43 ± 0.17 dB; pre-injection vs. post-early: P = 0.0; post-early vs. post-late: P = 5.90e-204; pre-injection vs. post-late: P = 0.0). **(D)** Song duration distributions for songs across different PAG TeLC experimental timepoints (pre-injection: n = 157 songs, 8.37 ± 0.20 s; post-early: n = 153 songs, 6.22 ± 0.15 s; post-late: n = 54 songs, 4.64 ± 0.20 s; pre-injection vs. post-early: P = 5.73e-16; post-early vs. post-late: P = 4.81e-8; pre-injection vs. post-late: P = 1.53e-19). **(E)** Distribution of the stop rate (*r*_min_)—one of the patterning parameters in our song rhythm model—for songs in (D) (pre-injection: 7.27 ± 0.16 Hz; post-early: 10.95 ± 0.17 Hz; post-late: 12.57 ± 0.25 Hz; pre-injection vs. post-early: P = 2.25e-33; post-early vs. post-late: P = 2.04e-6; pre-injection vs. post-late: P = 6.52e-24). **(F)** Song duration versus the stop rate for songs in (D). The black line represents the model prediction, based on a mean slope of -0.11 and a mean start rate of 19.15 Hz. **(G)** Distribution of the duration of the longest note for songs in (D) (pre-injection: 84.1 ± 1.8 ms; post-early: 48.3 ± 1.0 ms; post-late: 43.2 ± 2.0 ms; pre-injection vs. post-early: P = 4.18e-35; post-early vs. post-late: P = 0.001; post-early vs post-late: P = 2.76e-2). **(H)** Song duration distribution for intact males and females when alone or socially interacting in PAIRId (male: n = 370 songs from 6 mice, 6.28 ± 0.07 s; female: n = 86 songs from 4 mice, 3.58 ± 0.10 s; P = 6.65e-38). **(I)** Same as (E), but for songs in (H) (male: 8.27 ± 0.13 Hz; female 13.10 ± 0.30 Hz; P = 1.21e-30). **(J)** Same as (F), but for songs in (H). The black line represents the model prediction with a mean slope of -0.18 and a mean start rate of 24.23 Hz. **(K)** Same as (G), but for songs in (H) (male: 75.1 ± 1.1 ms; female: 49.2 ± 1.3 ms; P = 2.72e-24).

Given that the clPAG modulates song rhythm via the stop rate *r*_min_, we wondered if this mechanism might also account for natural variability in song duration. Consistent with previous work (15, 45), we observed songs are longer in males than in females (**Figure 6H**). Decomposing each song into its constituent parameters, we found that neither the start rate nor the slope significantly differed between the sexes (**Figure S4C**). However, stop rates were significantly lower for males compared to females (**Figure 6I**). Again, across all songs, durations closely followed the theoretical relationship with *r*_min_ predicted by our model. Thus, the longer songs in males reflect their ability to maintain the vocal pattern (frequency modulation) at lower stop rates—effectively extending the songs by adding more long notes at the end (**Figure 6K**). Therefore, the parameter most affected by clPAG silencing— the stop rate—also accounts for natural sexual dimorphism in song production. We conclude that clPAG is a key locus not only for orchestrating distinct vocal modes (songs and USVs), but also driving the graded, sexually-dimorphic natural variability in song production.

## Discussion

In this study, we leveraged the rich vocal behavior of the singing mouse (*Scotinomys teguina*) to investigate the organizational logic of multifunctional motor circuits. We developed a novel behavioral assay (PAIRId) that enables precise attribution of vocalizations to individual animals during social interactions solving a major technical bottleneck in bioacoustics (**Figure 1**). Using PAIRId, we found that singing mice employ two categorically distinct vocal modes: soft, unstructured USVs and loud, temporally patterned songs, often in quick succession during social encounters (**Figure 1**). We derived a simple linear model that captures the rhythmic structure of the songs with just three parameters (**Figure 2**). Despite their dramatic differences in acoustic properties, temporal organization, and social context, both songs and USVs share peripheral sound production mechanisms, vocal-respiratory coupling, and central neural control by the clPAG (**Figure 3, 4, 5**). Using our mathematical model, we demonstrated how progressive silencing of clPAG neurons systematically alters specific aspects of song production before eliminating all vocalizations (**Figure 6**). Collectively, our findings suggest a potential principle of multifunctional circuit reuse: a shared phonatory control pathway supports multiple vocal modes through amplitude modulation (AM) of individual notes and frequency modulation (FM) of global motor patterning (**Figure 7**). This work also offers a window into how neural circuits could be modified during evolution to diversify vocal repertoire in mammals, providing insights into the mechanistic basis of behavioral evolution.

**Fig. 7.**
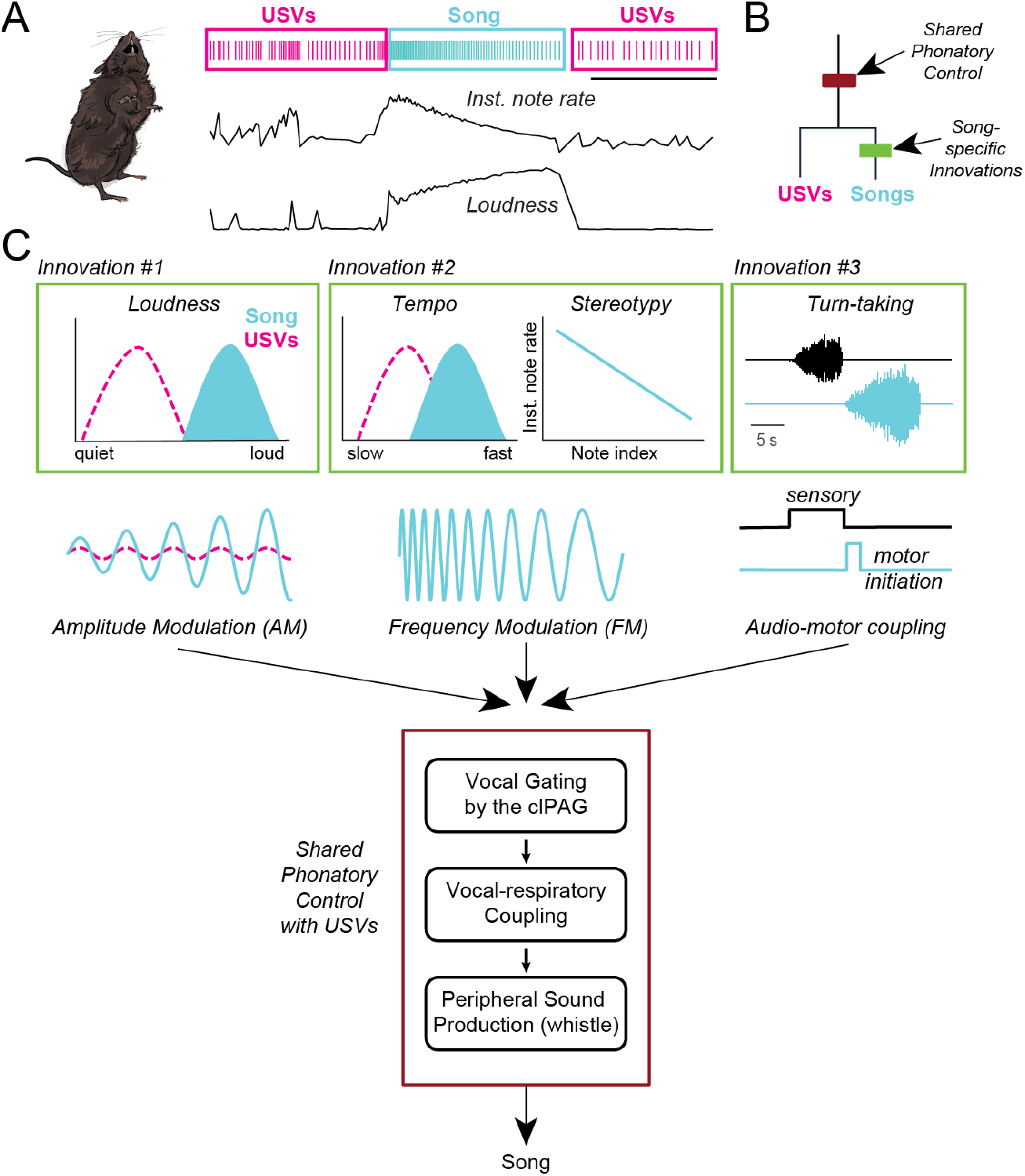
Working model of vocal motor control in the singing mouse. **(A)** Singing mice can produce two categorically distinct vocal modes in rapid succession–soft, unstructured USVs (magenta) and loud, patterned songs (cyan). **(B)** Songs and USVs share a phonatory control mechanism (maroon), while songs involve three separate innovations (green). **(C)** Cartoons depicting the three song innovations (green): (1) the notes are much louder than USVs, (2) songs include notes produced at much higher rates than USVs and arranged in highly stereotyped rhythms, and (3) as demonstrated previously, songs can be used in tightly coupled antiphonal turn-taking behavior known as countersinging. Together, these behavioral innovations imply that songs arise from specific modulations to existing phonatory pathway (maroon), including amplitude modulation for individual notes, frequency modulation for song rhythm, and audio-motor coupling for turn-taking.

### Two vocal modes: conserved USVs and novel songs

Our behavioral results suggest that singing mice have evolved a novel song mode while retaining ancestral USVs common to other rodents. The acoustic properties of singing mouse USVs—their soft amplitude, reduced temporal stereotypy, and usage during close-range social interactions—closely parallel USVs observed in laboratory mice (26, 46), despite differences in pitch that can be attributed to species-specific variations in laryngeal morphology (43, 44). The shared bio-physical mechanisms of sound production through aerodynamic whistles and vocal gating by the clPAG further support this homology at a mechanistic level (30, 38). This conservation is particularly notable given the phylogenetic distance between singing mice (Cricetidae) and laboratory mice (Muridae), suggesting that USV production represents a deeply conserved trait across multiple rodent families. Indeed, species producing USV-like vocalizations can be found across every studied subfamily within both Cricetidae and Muridae (28, 29, 34), providing strong evidence that USVs constitute an ancestral vocal mode that predates the divergence of these lineages approximately 20-25 million years ago (47).

In stark contrast, singing mouse songs represent a derived, novel vocal behavior unique to this lineage. Although songs share the same phonation mechanism with USVs, they differ significantly in their high amplitude, complex temporal patterning, and specialized usage in long-distance communication. This elaborate song mode may have evolved in response to a combination of selective pressures: long-distance communication in a diurnal cloud forest niche (14, 48), signaling costs linked to physiological conditions (49, 50), and sexual selection for display elaboration (51, 52). Similar songs, although not as elaborate, have been reported in a few other closely related species (*Baiomys* and *Scotinomys*) (15, 35). This restricted phylogenetic distribution suggests that songs are a recently evolved behavioral innovation (∼6.5 million years ago (47, 53)).

The stereotyped temporal progression of the songs can be captured by a simple linear model defined by three interpretable parameters. This compact parameterization not only accurately describes the structure of the behavior but also enables us to decompose variability into distinct components, each of which may be linked to separable neural substrates. In this way, the model provides a principled framework for investigating the neural mechanisms underlying specific aspects of song production. More broadly, singing mouse songs may exemplify a special class of natural behaviors— like birdsong (54), woodpecker drumming (55), or locomotor gaits (56)—that are highly stereotyped, rhythmic, and low-dimensional, making them particularly amenable to quantitative analysis. Therefore, the song system in the singing mouse offers an exciting opportunity for linking neural dynamics with motor output, allowing mechanistic circuit dissection of a natural behavior.

### Neural substrates of song production

The PAG plays a conserved role in the control of instinctive behaviors across vertebrates (57–62), with particular importance for vocalization (19, 20). Stimulation of the PAG (also called the “central gray”, and in birds, the “dorsomedial nucleus of the intercollicular complex”) elicits species-typical vocalizations across vertebrates, including fish (63), birds (64–66), bats (67), rodents (38, 68, 69), and primates (70–72). Further, bilateral PAG lesions cause mutism in both learned (speech) and innate vocalizations in humans (73), as well as mutism in many other species (63, 74–78).

Our findings in singing mice align with this body of evidence, confirming the critical role of clPAG in vocal control. However, our results extend the role of PAG beyond vocal gating by demonstrating its involvement in the moment-by-moment control of a complex, temporally patterned vocal behavior— singing. We demonstrate that the clPAG is essential for individual note phonation, high amplitude, and rhythmic structure over many seconds that together define song production. In particular, our mathematical model reveals how clPAG can implement a simple parametric control mechanism—the regulation of stop rates—that generate the natural variability in song duration observed both within and between sexes (15, 45).

While our study establishes the clPAG as a key locus in the song production pathway, future studies will elucidate the precise details of how neural activity within the clPAG generates songs. For example, although synaptic silencing of the clPAG eliminated both songs and USVs, optogenetic activation evoked only USVs. This dissociation is reminiscent of findings in songbirds, where experimental stimulation of motor pathways can elicit individual syllables but not complete songs (64, 65, 79, 80), and may reflect that the intrinsic neural dynamics required for song production are more complex than the simple vocal gating sufficient for USVs. Extending beyond the PAG, the flexible yet tractable vocal behaviors of the singing mouse provide a valuable opportunity for investigating hierarchical vocal motor control from the cortex to the brainstem in a mammal (17, 18, 81, 82).

### The PAG as a key locus for rapid evolutionary diversification

Evolutionary modification of ancestral neural circuits provides a pathway for rapid behavioral diversification. The midbrain PAG is anatomically well positioned to receive ethologically relevant contextual inputs from the forebrain and to coordinate diverse motor output programs (83–85), leading to a recent proposal positing the PAG as a key driver in the evolution of innate motor behaviors across species (86). We put this idea to test and indeed identified the clPAG as a critical locus for the evolutionary diversification of vocal behavior, supporting both the ancestral USVs and the derived song. The song circuit likely not only co-opts the PAG’s core phonation motor program used for USVs but also redeploys it into the distinct contextual inputs that drive the differential social use of these two vocal modes. While much remains to be done, the coexistence of the ancestral USVs and the novel songs in the same animal often produced in rapid succession offers a powerful model for understanding how changes in neural circuits drive novel behaviors.

Neural circuit co-option may represent a common strategy for evolving complex behaviors. Rather than evolving separate circuits *de novo*—a process requiring coordinated changes across multiple organizational levels—evolution appears to favor modifying ancestral circuits to operate in new regimes (7, 57, 87–92). Beyond vocal communication in singing mice, quantitative modifications of preexisting circuit nodes is widespread at the molecular, genetic, and anatomical levels, representing a general principle for the rapid evolutionary emergence of biological innovations across scales and systems.

## RESOURCE AVAILABILITY

### Lead contact

Requests for further information and resources should be directed to and will be fulfilled by the lead contact, Arkarup Banerjee (*abanerjee@cshl*.*edu*).

### Materials availability

This study did not generate new unique reagents.

### Data and code availability

All data and code are available from the lead contact upon request.

## ACKNOWLEDGMENTS

We are grateful to Priyanka Gupta and Devon Cowan for performing the thermistor implantation surgery and technical support. Luke Bemish contributed to earlier iterations of the song rhythm model. Bo Li generously provided the Flex TeLC virus. We thank Florin Albeanu, Benjamin Cowley, Priyanka Gupta, Michael Long, Stephen Shea, and members of the Banerjee lab for their helpful comments on earlier versions of the manuscript. We thank the staff of the Cold Spring Harbor Laboratory Animal Resource for their dedicated animal care. We thank the creators and maintainers of the open-source software used in this project. We also thank Taylor Sterry for visual illustrations of the mice and the PAIRId apparatus.

## AUTHOR CONTRIBUTIONS

*Conceptualization*: AB, XMZ, CEH

*Methodology* : XMZ, CEH, AB

*Investigation*: XMZ, CEH, MBD

*Data Curation*: CEH, XMZ

*Software*: XMZ

*Formal Analysis*: XMZ, CEH

*Visualization*: XMZ, CEH, AB

*Writing—original draft* : AB

*Writing—review & editing*: XMZ, CEH, AB

*Supervision*: AB

*Project Administration*: AB

*Funding Acquisition*: AB

## DECLARATION OF GENERATIVE AI AND AI-ASSISTED TECHNOLOGIES IN THE WRITING PROCESS

The authors used ChatGPT and Claude to improve the readability of the manuscript. The authors take full responsibility for the content of the published article.

---

**Fig. S1.**
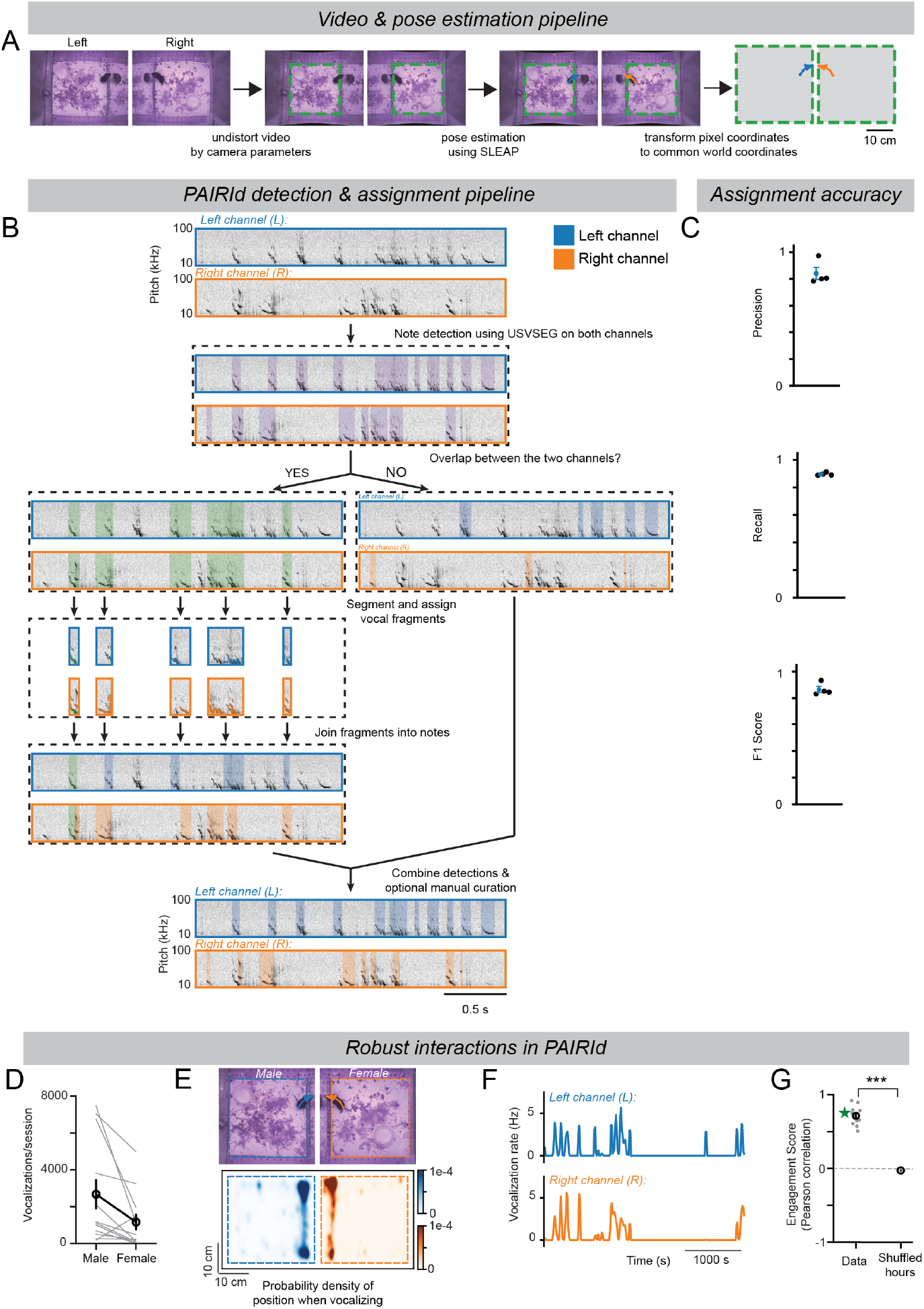
Data processing pipelines and vocal interaction quantification of PAIRId, related to Figure 1. **(A)** The PAIRId video processing pipeline aligns two mice in common world coordinates. In steps, the distortion produced by the lens is removed relative to the enclosure floor, followed by pose estimation using SLEAP. The output positions of the 5-node skeleton are then transformed from pixel values to common world coordinates. **(B)** Schematic overview of the PAIRId audio processing pipeline to attribute vocalizations to individual mice. **(C)** Accuracy performance metrics of PAIRId assignment compared with “ground truth” produced via human manual annotation. False positives for a given channel are defined as a PAIRId detection that lacks a match in ground truth detections for either the given channel or unassigned. False negatives for a given channel are defined as a ground truth detection that lacks a match in PAIRId detections for either the given channel or unassigned. *Top*: Precision, or the amount of true positives divided by the total of true positives and false positives, is 0.84 ± 0.04. *Center* : Recall, or the amount of true positives divided by the total of true positives and false negatives, is 0.89 ± 0.01. *Bottom*: F1 score, the harmonic mean of precision and recall as a combined performance metric, is 0.86 ± 0.02. For each, black circles are the values of the four ground truth hours compared with PAIRId algorithm, and blue circles are the mean ± sem for these four values. **(D)** Number of vocalizations in PAIRId by the male (2673.3 ± 818.0, n = 12 sessions of 5 hours each, 6 animals) and the female (1162.3 ± 451.0, n = 12 sessions of 5 hours each, 4 animals). **(E)** Density of mouse locations (centroid) during vocalizations (n = 34,877 notes in 11 hours of high vocal engagement from 4 males and 3 females) **(F)** Rates of vocal production of two interacting mice (*top*: male; *bottom*: female) in an example hour with high vocal engagement. Epochs of vocal production were correlated between the two animals (Pearson correlation = 0.75). **(G)** Summary of Pearson correlations between the vocal rates of interacting mice during periods of high vocal engagement. The correlation coefficient for data (0.71 ± 0.04, n = 11 hours) is significantly higher than that of the shuffled control (P = 5.03e-8). Green star indicates the example hour in (F). Unless stated otherwise, values reported are Mean ± SEM and hypothesis testing was performed using Mann-Whitney U test.

**Fig. S2.**
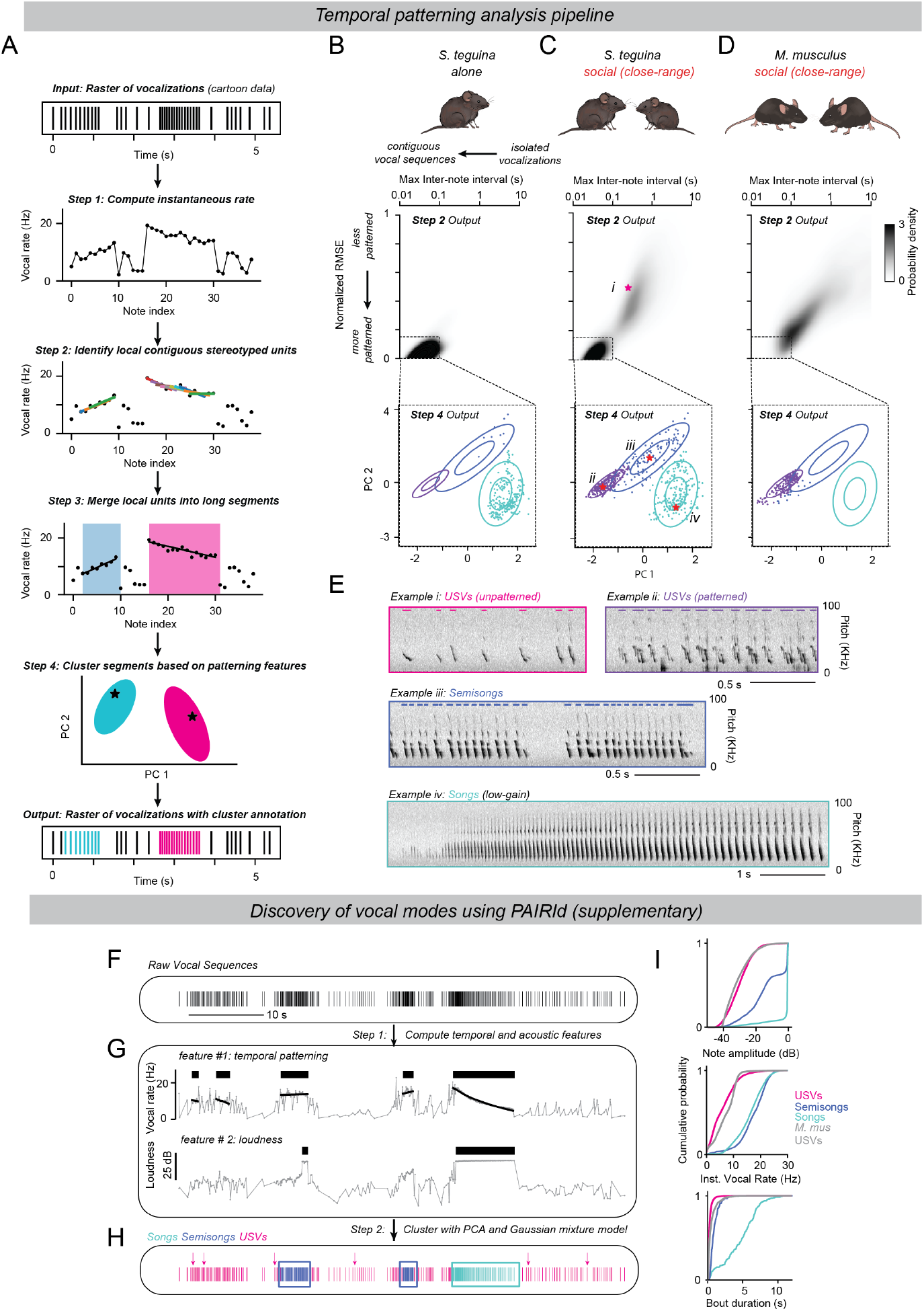
Temporal patterning analysis in the identification of distinct vocal modes, related to Figure 1. **(A)** Schematic overview of the analysis pipeline for identifying stereotyped vocal sequences, using cartoon vocalizations as a demonstration. (*1*) First, instantaneous note rate of vocalizations—the inverse of the difference between two consecutive note-start times—was computed and plotted against note index. (*2*) Then, using a sliding window of notes, local contiguous units with stereotypy are identified (colored lines). A window is deemed temporally patterned if it exhibits both temporal contiguity (via a low max inter-note interval), and stereotyped note sequencing (via a low NRMSE). (*3*) Consecutive windows (overlapping and/or with 0.5 s maximum gap separation) meeting these criteria are merged into longer segments (blue and red highlights). Merged segments are characterized via RANSAC regression to robustly fit a linear model (black lines). (*4*) Key patterning features of the linear model (maximum rate, minimum rate, and number of notes from RANSAC) are used for clustering the patterned segments with a Gaussian mixed model (GMM) on principal components. In this cartoon, the blue segment falls into the cyan cluster and the pink segment into the magenta cluster. (*output*) Notes from the identified segments are assigned a category based on the clusters from the temporal patterning analysis (cyan, magenta), while others are considered unpatterned (black). **(B-D)** Intermediate steps from the temporal patterning analysis of all vocalizations of the listed conditions: (B) singing mice alone, (C) singing mice with nearby conspecifics in PAIRId, and (D) laboratory mice in a nearby social setting. The top plots are the values for all windows from step 2, and dotted lines the thresholds for patterning and contiguity. The bottom plots are the values of the principal components of the features of linear models of patterned, merged segments. Ellipses represent the first and second standard deviations of the clusters found via GMM. **(E)** Representative example spectrograms of vocalizations produced by singing mice of the 4 clusters—(i) unpatterned USVs, (ii) patterned USVs, (iii) semisongs, and (iv) songs. Overlaid colored lines indicate vocalizations of the focal mouse. The temporal features of examples (i)–(iv) are denoted with stars in (C). **(F)** Raster plot illustrating vocalizations of a singing mouse in the PAIRId paradigm during an example 60-second period, with each line marking the onset of a vocalization. **(G)** Vocal sequences were characterized by substantial variation in temporal stereotypy (top) and loudness (bottom). *Top*: Temporal patterning features were extracted by the procedure outlined in (A) with the black bar indicating long, stereotyped vocal sequences. *Bottom*: Amplitude features were computed for the vocalizations, with the black bar highlighting those with high loudness. **(H)** The same raster plot as in (F), with lines color-coded by the vocal modes (songs, cyan; semisongs, blue; USVs, magenta) identified by combining temporal stereotypy and loudness features. **(I)** *Top*: Cumulative distribution of note loudness for each vocal mode (singing mouse USV: n = 26,246 notes from 10 mice, -29.78 ± 0.05 dB; singing mouse semisong: n = 2,100 notes from 9 mice, -13.06 ± 0.25 dB; singing mouse song: 17,681 notes from 10 mice, -2.27 ± 0.05 dB; laboratory mouse USV: n = 7,295 notes from 3 mice, -30.82 ± 0.09 dB). *Center* : Cumulative distribution of instantaneous note rate (singing mouse USV: 6.26 ± 0.03 Hz; singing mouse semisong: 17.24 ± 0.11 Hz; singing mouse song: 16.16 ± 0.04 Hz; laboratory mouse USV: 7.24 ± 0.05 Hz). *Bottom*: Cumulative distribution of vocal bout duration (singing mouse USV: 0.31 ± 0.00 s; singing mouse semisong: 0.96 ± 0.05 s; singing mouse song: 4.64 ± 0.15 s; laboratory mouse USV: 0.56 ± 0.02 s).

**Fig. S3.**
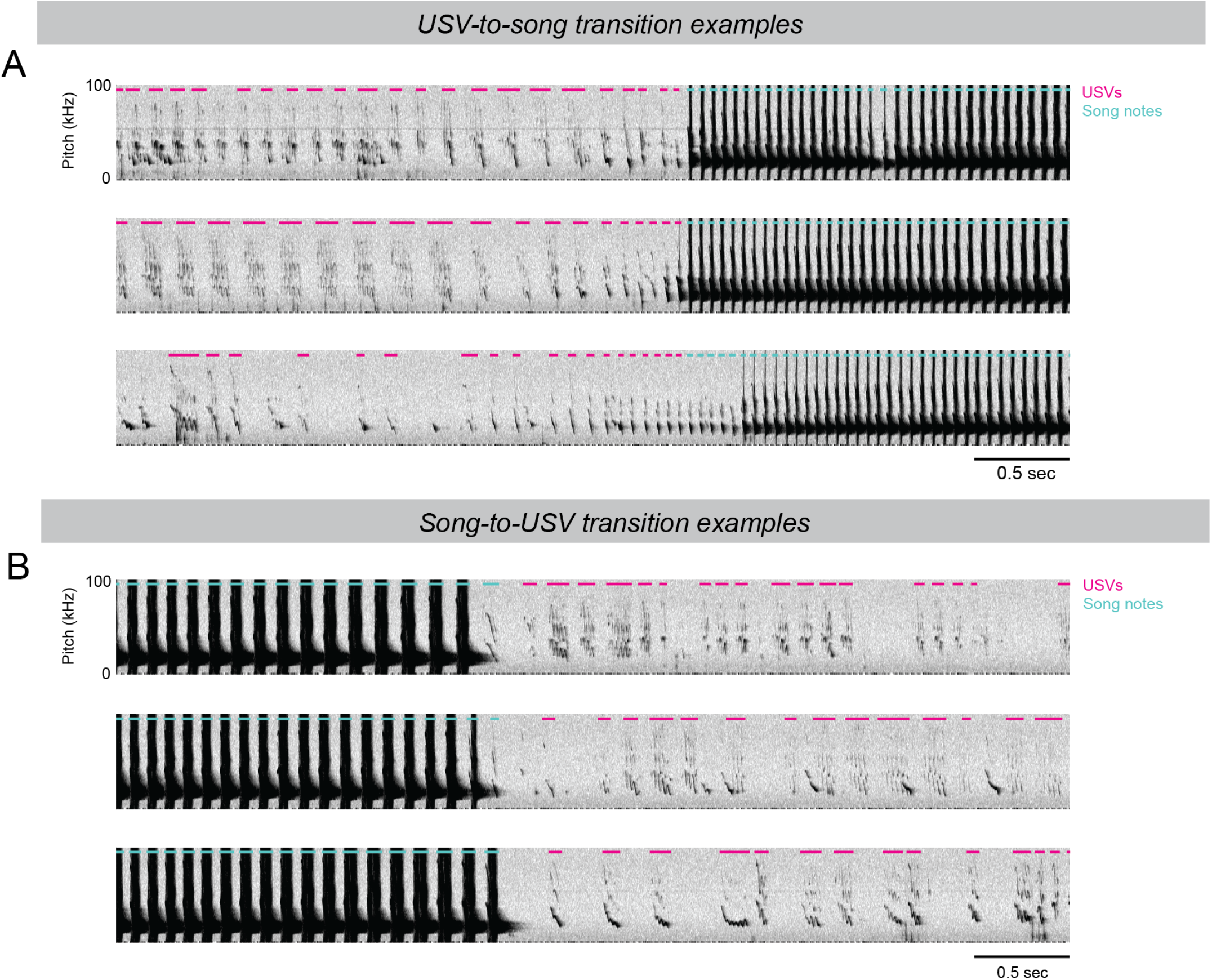
Rapid, smooth transitions between USVs and songs, related to Figure 1. **(A)** Representative example spectrograms displaying a singing mouse transitioning from USVs to songs. **(B)** Representative example spectrograms displaying a singing mouse transitioning from the final notes of a song to USVs. Overlaid colored lines indicate vocalizations of the focal mouse, categorized as USV (magenta) or song (cyan).

**Fig. S4.**
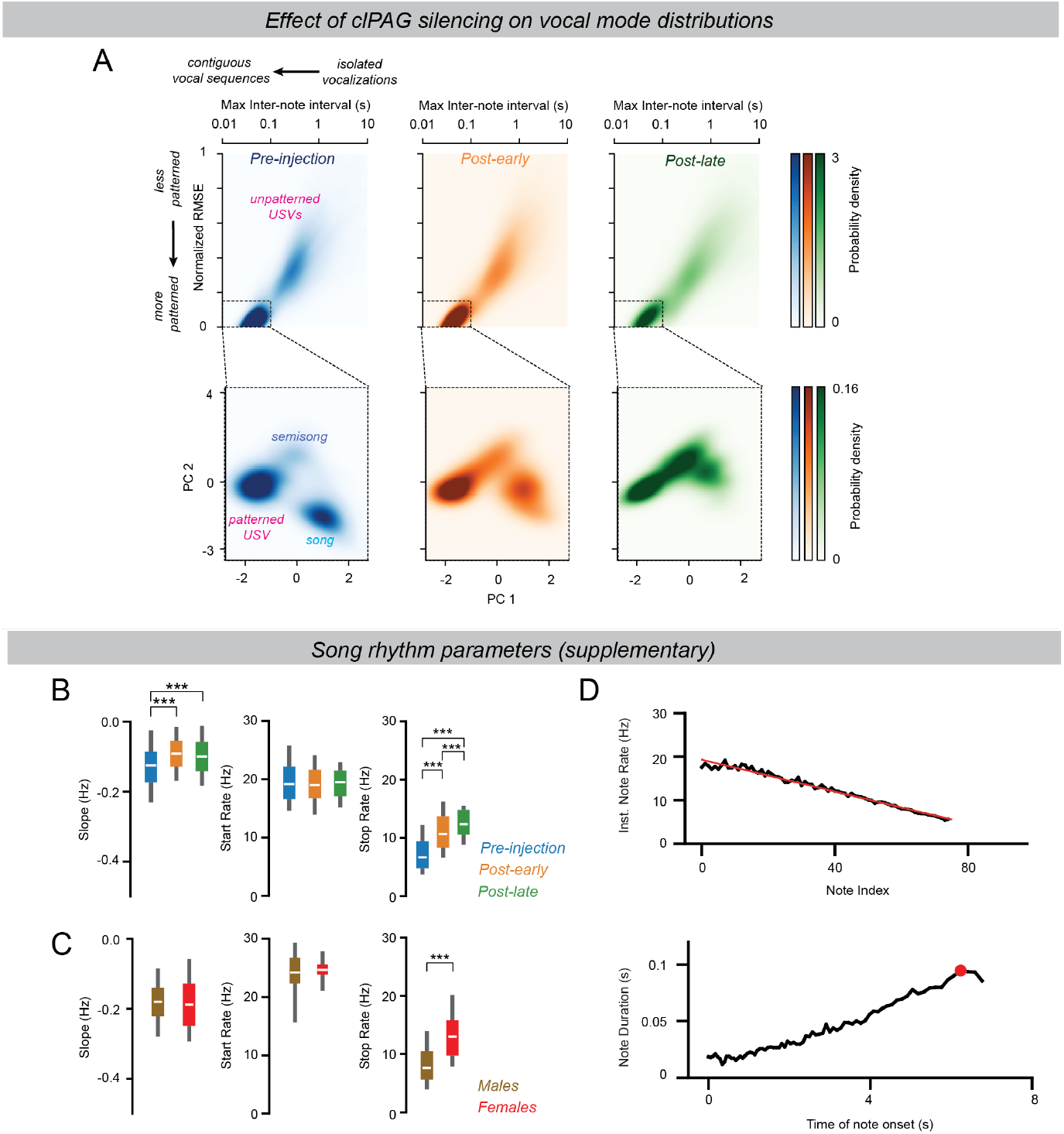
Effects of TeLC-mediated silencing of the clPAG on the vocal Repertoire, with a focus on songs, related to Figures 5–6. **(A)** Intermediate steps from the temporal patterning analysis of all vocalizations before the TeLC virus injection in the clPAG (left; n = 428 segments from 39,596 notes), during the post-early period (middle; n = 408 segments from 34,826 notes), and during the post-late period (right; n = 252 segments from 21,084 notes). The top panels show the stereotypy and contiguity of local units, and the bottom panels show the temporal features of the merged long segments. **(B)** Distribution of song patterning parameters of songs before TeLC virus injection in the clPAG (n = 157 songs), during the post-early period (n = 153 songs), and during the post-late period (n = 54 songs). *Left*: slope (pre-injection: -0.13 ± 0.00; post-early: -0.09 ± 0.00; post-late: -0.09 ± 0.01; pre-injection vs. post-early: P = 1.15e-14; post-early vs. post-late: P > 0.05; post-early vs. post-late: P = 1.43e-5). *Middle*: start rate (pre-injection: 19.38 ± 0.18 Hz; post-early: 18.97 ± 0.18 Hz; post-late: 19.00 ± 0.29 Hz; pre-injection vs. post-early: P > 0.05; post-early vs. post-late: P > 0.05; post-early vs. post-late: P > 0.05). *Right*: Stop rate (pre-injection: 7.27 ± 0.16 Hz; post-early: 10.95 ± 0.17 Hz; post-late: 12.57 ± 0.25 Hz; pre-injection vs. post-early: P = 2.25e-33; post-early vs. post-late: P = 2.04e-6; pre-injection vs. post-late: P = 6.52e-24). **(C)** Distribution of patterning parameters of songs in males and females. *Left*: slope (male: -0.18 ± 0.00; female: -0.19 ± 0.01; P > 0.05); *Middle*: start rate (male: 24.19 ± 0.15 Hz; female: 24.39 ± 0.19 Hz; P > 0.05). *Right*: stop rate (male: 8.27 ± 0.13 Hz; female 13.10 ± 0.30 Hz; P = 1.21e-30). **(D)** Trajectories of an example song visualized in two ways. *Top*: Instantaneous note rate is plotted against note index, same as in our model of the song rhythm. *Bottom*: Note duration is plotted against note onset time. The red circle highlights the longest note in the song.

**Fig. S5.**
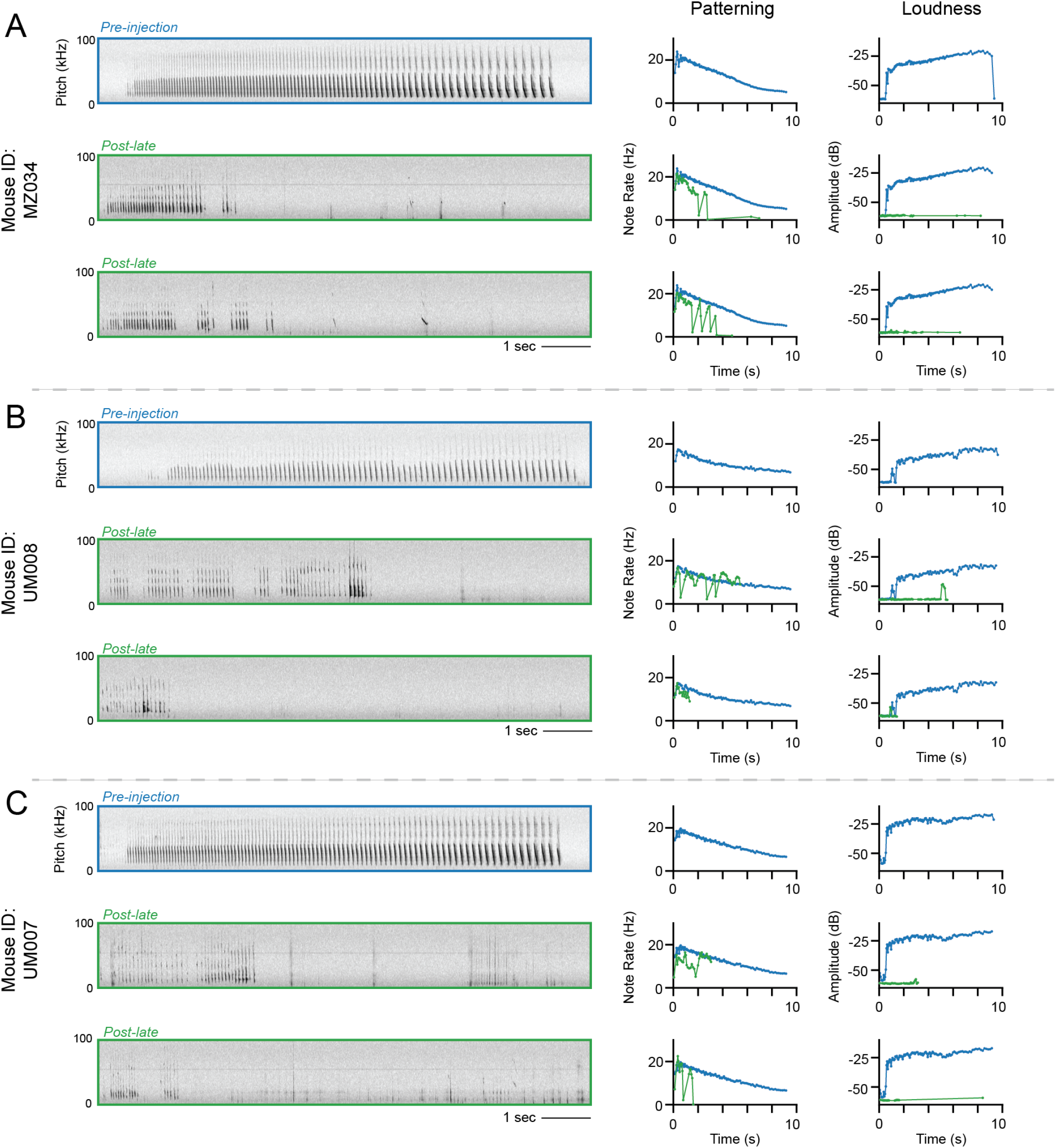
Examples of final song-like sequences by TeLC-mediated silencing of the clPAG, related to Figures 5–6. **(A-C)** Examples of the final song-like sequences wherein multiple song properties are heavily degraded as a result of TeLC expression in the clPAG. Each triplet represents a different mouse: the top panel shows a representative pre-injection song with blue-outlined spectrograms and corresponding patterning and loudness traces; the subsequent panels display two final song-like sequences from the post-late period (in green), with the pre-injection traces provided as a reference.

**Supplementary Video 1 A song of an Alston’s singing mouse (*S. teguina*), related to Figure 1 and 2**. Top: video recording of an adult male mouse singing. Bottom: spectrogram of the song, illustrating the temporal patterning of note progression.

**Supplementary Video 2 Two singing mice vocalizing in the PAIRId paradigm, related to Figure 1**. Top left: video recordings of each enclosure. Top right: the poses of both mice in common world coordinates. Bottom: spectrograms from each enclosure, overlaid with vocalizations detected and attributed to individual mice. The pitch of high-gain audio was reduced by a factor of eight for human hearing range.

**Supplementary Video 3 Optogenetic activation of the clPAG in a singing mouse elicited USVs, related to Figure 4**. Top: Video recording of the mouse, displaying the 4-second tonic 473 nm laser stimulation. Bottom: Spectrogram of the mouse’s vocalizations, demonstrating USVs throughout the stimulation period. The pitch of high-gain audio was reduced by a factor of eight for human hearing range.

## Materials & Methods

### Experimental Model

#### Animal statement

All animal care and experiments were conducted according to protocols approved by the Cold Spring Harbor Laboratory Institutional Animal Care and Use Committee, and comply with the National Institutes of Health Guide for the Care and Use of Laboratory Animals.

#### Animals

Adult, laboratory-reared, outbred male and female Alston’s singing mice (*Scotinomys teguina*), aged 3–20 months, were selected from the colony maintained at Cold Spring Harbor Laboratory. This colony originated from the New York University Langone Medical Center colony (1), which was descended from wild-captured *S. teguina* from La Carpintera and San Gerardo de Dota, Costa Rica. The sex of the singing mice was confirmed at weaning by genotyping the Y-chromosome’s *Sry* gene, using the *Zfy-Zfx* genes as a positive control (2). Singing mice were kept at 20–22°C under a 12:12-hour light-dark cycle. They were housed in Thoren Systems #8 enclosures (30.80 × 40.60 × 22.23 cm; Worcester, MA) with corn cob bedding and enrichment items, including sphagnum moss (Galapagos Pet, Santa Barbara, CA), a running wheel (InnoDome + InnoWheel; Bio-Serv, Flemington, NJ), a paper hut (Bio-Hut; Bio-Serv), and a red transparent polycarbonate tube (10 cm × 5 cm, Mouse Tunnel; Bio-Serv). The singing mice were provided with food (a 1:1 mixture of Purina Cat Chow and Mazuri Exotic Animal Nutrition Insectivore pellets) and water *ad libitum*, supplemented with dried mealworms. In a different room from singing mice, C57Bl/6J laboratory mice were kept at 20–22°C in Thoren Systems #9 enclosures (19.56 × 30.91 × 13.34 cm) with corn cob bedding under a 12:12-hour light-dark cycle. Water and mouse chow were available *ad libitum*.

### Experimental Procedures

#### PAIRId social dyad assay

To capture the vocal repertoire of rodents when they are close to one another and reliably identify which individual in a dyad produced each vocalization, we designed a “two-enclosure” behavioral assay which leverages partial acoustic isolation and two microphones (Figure 1). We refer to this assay as “PAIRId” (**p**artial **a**coustic **i**solation **r**eveals **id**entity). The enclosures were custom-built transparent acrylic rectangular boxes (outer dimensions: 12 × 12 × 18 inches, wall thickness: 0.25 inches; shopPopDisplays, Woodland Park, NJ). One face of each box was drilled with nine 0.25-inch holes using cutting drill bits: five holes positioned 2 inches above the inner floor, and four additional holes offset from the top row, 1.5 inches above the inner floor. A removable floor, covered with AlphaPad bedding (Shepherd Specialty Papers, Watertown, TN), was inserted into each enclosure to facilitate cleaning between sessions. Inside of a controlled acoustic environment box, we placed an aluminum breadboard designed to fit two of these enclosures and maintain a .932 cm gap between the outer dimensions of each. Considering the thickness of the two inner walls, the effective division between two rodents is 2.2 cm. The perforated faces of the acrylic boxes were facing the other across this gap, allowing the mice to acoustically interact with one another. To ensure proper ventilation, we fed external air into each box via tubing. To the breadboard we also attached two custom assembled cranes such that a microphone and a video camera could be lowered into each enclosure to the consistent height of 12 in between trials (crane materials from 80/20 Inc., Columbia City, IN). A third microphone was placed in the chamber and set to low gain to record the enclosure. Finally, when each enclosure contained their rodent, microphone, and camera, we inserted custom-cut 2-inch-thick polyimide acoustic foam into the opening (Soundfoam HTC, The Soundcoat Company, Deer Park, NY). The foam was cut to the outer perimeter of the acrylic box and lightly compressed to fit the inner perimeter, creating acoustic dampening between the enclosures. Each enclosure contained an Avisoft CM16/CMPA microphone, with a third microphone positioned outside the two enclosures. All three microphones were powered and recorded by an Avisoft UltraSoundGate 416H device, saving WAV files at a 250 kHz sampling rate using Avisoft-RECORDER software. Each enclosure also housed a FLIR Blackfly USB camera (BFS-U3-20S4C-C, Teledyne FLIR, LLC) equipped with a 4.5 mm fixed-focal-length lens (C Series #86-900, Edmund Optics). Frames were captured at 50 frames per second, triggered by a custom-programmed Arduino Mega 2560 R3. The same Arduino controlled the start and stop triggers for audio recording, ensuring synchronized audio and video.

#### “Alone” assay

To capture the repertoire of a singing mouse in isolation while accounting for the novelty of the PAIRId enclosure, we used clear acrylic boxes with the same footprint as the PAIRId enclosure but with a shorter height (10 in) to fit inside soundproof acoustic-foam-lined MedAssociates cabinets (Fairfax, VT). Each box had a wire lid and was equipped with both high- and low-gain microphones outside but near the box. Audio was recorded using an Avisoft UltraSoundGate 416H device, saving WAV files at a 250 kHz sampling rate via Avisoft-RECORDER software.

#### Alone-social comparison experimental timeline

In two cohorts of 3 males and 2 females each (total of 6 males and 4 females aged 4-11 months), each combination of opposite sex dyads was subjected to the following experimental timeline: On Day 0, each singing mouse was removed from its home enclosure and placed in a short acrylic “alone” box to acclimate overnight. On Day 1, the singing mice were recorded individually in the alone boxes for 5 hours. After recording, they were transferred to the PAIRId enclosures, where they acclimated to the experimental setup in acoustic isolation from one another overnight. On Day 2, at approximately the same time, the two PAIRId enclosures were placed next to each other in the PAIRId setup, allowing the singing mice to interact while being recorded for 5 hours as described above. Following this, the session was complete, and the singing mice were transferred back to their home cages. All 12 combinations of opposite sex dyads sessions were recorded over the course of nine days on a schedule that allowed at least 24 hours of time spent in home cage between sessions for individual singing mice. This design resulted in a dataset with each male represented in two sessions, one with each female of its cohort, and each female in three sessions, one with each male.

#### Laboratory mouse dyad experiment

To compare acoustic parameters of lab mice with those measured in singing mice, we re-analyzed the *Mus musculus* male-female dyad dataset from (3). In this dataset, a cohort was selected of three male and three female C57Bl/6J mice aged ∼two months old (58-70 days) such that the sexes were not littermates. All mice were singly housed and isolated for at least nine days before social exposure to increase the probability of vocalization (4). In advance of recording, the female mouse was placed into a clean cage (Thoren Systems #8, Worcester, MA; 30.80 × 40.60 × 22.23 cm) lined with clean Alpha-pad cotton paper. After a period of acclimatization (min: 30 mins, max: 12 hours), a male was introduced to the cage with the female. Audio of the pair was recorded for 1 hour using two Avisoft UltraSoundGate 116H devices and two Avisoft CM16/CMPA microphones (with high and low gains), synchronized via a custom external trigger, with WAV files written with a 250 kHz sampling rate using Avisoft-RECORDER software. Female laboratory mice rarely vocalize in male-female dyads during social interactions of this length (5, 6). Consistent with this, visual inspection of spectrograms revealed no apparent instances of overlapping USVs that would indicate that the male and female mouse were vocalizing at the same time.

#### Surgical Procedures

Mice subjected to surgery were placed into an induction chamber with 1-2% isoflurane. Hair was clipped from the surgical site. The mouse was then placed onto a heating pad on a stereotaxic instrument (Kopf model 940, Tujunga, CA). The mouse’s front teeth were latched onto a bite bar, the head secured with non-rupturing ear bars, and the head levelled. Nonsteroidal anti-inflammatory drug meloxicam was administered subcutaneously at 5 mg/kg. Following these preparations, the mouse was subjected to either 1) implantation of a thermistor, 2) injection of viral vector followed by implantation of an optogenetic fiber, or 3) injection of viral vectors.

#### Thermistor implantation

Chronic implantation of an intranasal thermistor is a well-established method for estimating respiration in rodents (7). As obligatory nose-breathers, a rodent’s nasal cavity warms during exhalation and cools when room-temperature air is inhaled due to the difference of a rodent’s internal temperature and experimental conditions. We implanted thermistors, adapted from McAfee et al 2016. Briefly, after preparation (see above), a midline incision was made over the skull, to the anterior edge of the nasal bone. A cavity for the thermistor was opened by drilling the nasal bone (A/P 3.1mm, M/L 0.5mm from nasal suture), and the thermistor implanted within. The thermistor was sealed using Kwik-Cast silicone sealant, and the implant was secured to the skull with layers of Vitrebond, Metabond, and dental acrylic.

#### Stereotaxic viral injection & optogenetic cannula implantation

Adeno-associated viruses (AAVs) originally developed for laboratory mice also infect neurons and express their packaged cargo in singing mice. In this study, we leverage this to 1) activate neurons optogenetically using channelrhodopsin and 2) silence neuronal synaptic transmission using tetanus toxin light-chain (TeLC). We first identified the caudolateral PAG region of singing mice as 4.2 mm posterior and 0.6 mm lateral relative to bregma and 2.3 mm ventral from the brain’s surface. Craniotomies were made using a dental handpiece and an FG ¼ carbide burr (Dentsply Sirona Midwest Tradition TL, Dentsply Sirona, Charlotte, NC). Viruses were injected using a Nanoject III (Drummond Scientific) at 2 nL/cycle with a 10 second interval. For the TeLC silencing experiments, five male singing mice (5-12 months old) were injected bilaterally in the clPAG with 80 nL of a 1:1 mixture of AAV2/DJ-hSyn-flex-TeLC-eYFP (Addgene #135391, custom packaged by WZ Biosciences) and AAV2/9-pENN.AAV.CamKII 0.4.Cre.SV40 (Addgene #105558). For the optogenetic activation experiments, four male singing mice (6-11 months old) were injected unilaterally in the clPAG with 150 nL of a 1:1:2 mixture of AAV2/9-EF1α double-floxed-ChR2-mCherry (Addgene #20297), AAV2/9-pENN.AAV.CamKII 0.4.Cre.SV40 (Addgene #105558) and sterile saline before fiber implantation. In the same surgery following this injection, an optogenetic cannula with a tapered tip (Optogenix, .39/200, active length 0.5mm, implant length 3mm, cLCF) was implanted and secured to the skull using Metabond. Subsequently, a headbar was also implanted and secured using Metabond, and the entire implant was protected by dental acrylic.

#### Histology

Mice were transcardially perfused with PBS followed by 4% paraformaldehyde (PFA), after which brains were dissected and post-fixed in 4% PFA overnight before being stored in PBS. Brains were sectioned into 100 µm coronal slices using a vibratome. To visualize tissue structure, select slices were stained with NeuroTrace 435/455 (Thermo Fisher Scientific, N21479) at a 1:30 dilution following the manufacturer’s protocol. Stained slices were mounted on glass slides using ProLong Gold Antifade mounting medium (Thermo Fisher Scientific, P36930) and imaged with an epifluorescence microscope.

#### Vocal-respiratory coordination experiment

To estimate a singing mouse’s respiration while vocalizing its full vocal repertoire, we implanted an intranasal thermistor. We used a muted female singing mouse (described below) as a stimulus to elicit the male’s vocalizations, ensuring all recorded vocalizations originated from the implanted male. During an experiment, the thermistor of the implanted mouse was first connected to an overhead rotary joint (Adafruit, #736) and then routed into a custom-built amplifier circuit. The implanted male was recorded alone or with the presence of a singing mouse in a custom transparent acrylic cylindrical enclosure (12 inch diameter, 12 inch tall). The amplified thermistor signal was recorded via an Intan RHD 1024ch Recording Controller (Intan Technologies, Los Angeles, CA). Vocalizations were captured using two Avisoft UltraSoundGate 116H devices and two Avisoft CM16/CMPA microphones, synchronized via a custom external trigger, with WAV files written with a 250 kHz sampling rate using Avisoft-RECORDER software. Synchrony with the thermistor signal was achieved via recording a copy of the trigger-on signal with the Intan RHD recorder.

#### Laryngeal phonation mechanism experiment

To determine the laryngeal phonation mechanism of the singing mice vocal repertoire, we recorded vocalizations from four male-female dyads (7-20 months, older singing mice freeze less and vocalize quickly after disturbance by an experimenter) in both air and heliox (80% He, 20% O_2_). Each dyad was placed in a Thoren Systems #8 enclosure with a removable acrylic floor covered with clean AlphaPad bedding, beneath which a perforated clear PVC tube connected to the heliox tank was positioned. The enclosure was housed inside a MedAssociates cabinet lined with acoustic foam to facilitate heliox accumulation and acoustic isolation. The singing mice were allowed to vocalize for 45–60 minutes before heliox was introduced at a flow rate of 5 L/min, for an additional 45-60 minutes. Vocalizations were captured using two Avisoft UltraSoundGate 116H devices and two Avisoft CM16/CMPA microphones, synchronized via a custom external trigger, with WAV files written with a 250 kHz sampling rate using Avisoft-RECORDER software.

#### Optogenetic activation experiment

Two weeks following the surgery, clPAG neurons were optogenetically activated with blue light from a 473 nm laser with tonic light stimulation for 1, 2, or 4 seconds. Vocalizations were captured using two Avisoft UltraSoundGate 116H devices and two Avisoft CM16/CMPA microphones, synchronized via a custom external trigger, with WAV files written with a 250 kHz sampling rate using Avisoft-RECORDER software. Copies of the external synchronization trigger signal and the laser signal were recorded on an Intan RHD 1024ch Recording Controller (Intan Technologies, Los Angeles, CA) for synchronization.

#### Tetanus toxin light chain inactivation experiment

In each experiment, a male-female singing mouse dyad was allowed to interact in the PAIRId setup for 24 hours as a “pre-injection” time point. After the baseline recording, the male of each dyad was subjected to injection of a virus mixture to silence neurons in the clPAG via tetanus toxin light chain. Following the injection and recovery from anesthetic on a heating pad (usually within a half hour), the perturbed singing mouse male was placed back into the PAIRId assay with its stimulus female and continuously recorded for four or five days.

### Analysis

#### Detection of vocalizations

To analyze vocalizations, we first segmented biotic sounds from silence in audio files, a necessary step for vocal analysis in both PAIRId and other paradigms. We used a modified version of USVSEG software (usvseg09r2) (8), which implements a signal processing algorithm for the detection of typically quiet rodent sounds from background noise. Briefly, USVSEG extracts vocalization events by generating a stable spectrogram using the multitaper method, flattening it in the cepstral domain to remove noise, applying thresholding, and estimating onset/offset boundaries. This robust, species-agnostic software allowed us to adjust parameters to suit the acoustic profiles of singing mice specifically. We modified the open-source software slightly: to improve inter-file consistency, we adjusted the threshold calculation for detecting biotic sounds from a noise-based standard deviation per file to a fixed value optimized per setup. USVSEG performed well for detecting quiet vocalizations; however, the loud vocalizations emitted by singing mice were more reliably segmented using a custom Python-based method optimized to handle reverberations. This method segmented loud notes based on the signal-to-noise ratio in acoustic power, calculated from a spectrogram. A rolling estimate of background noise was used to dynamically adjust the noise threshold for segmentation. The detections herein were merged with those of USVSEG, with redundancies removed by giving priority to the detected loud notes. For data recorded in the PAIRId social assay, segmentation was performed for the left and right microphones separately before assignment. For the “alone”, heliox, thermistor, optogenetic activation, and lab mouse dyad experiments that did not require individual assignment, these data would next be curated.

#### Assignment of vocalizations in the PAIRId setup

To determine the source of each vocalization in the PAIRId social assay, we compared detections from both enclosures. If a detection occurred on one side without an overlapping detection on the other, the source was assigned to that side. For overlapping detections, we distinguished between simultaneous vocalizations (“coincidence”) and acoustic bleed-through. In cases of coincidence, the spectro-temporal shapes of the vocalizations differ between channels; in contrast, similar shapes indicate bleed-through. To extract the spectro-temporal shapes, we segmented the vocal fragments on each side by thresholding the spectrograms. If a fragment appeared in only one channel (consistent with coincidence), that channel was assigned as the source of that fragment. If the fragment appeared in both channels (consistent with bleed-through), the source was assigned to the louder channel or marked as unknown if there was little difference in acoustic power. After assigning the vocal fragments, we joined fragments on each side into notes respectively. Finally, we combined the non-overlapping notes and notes assigned from overlaps to produce the final output. See Figure S1B for a visual overview.

#### Quantification of the performance of PAIRId setup

We first quantified the performance of the PAIRId setup by evaluating its assignment step. The final output combines non-overlapping notes—which reflect the hardware’s partial acoustic isolation performance—with notes assigned from overlapping detections using the algorithm. For each session, we calculated both the percentage of notes assigned solely by the hardware and the percentage of total notes assigned (Figure 1E). Next, we evaluated the full analysis pipeline by benchmarking it against four manually-annotated “ground truth” hours. During these sessions, we assessed the pipeline’s performance by quantifying true positives, false positives, and false negatives. For instance, a false positive on the left was defined as an assignment on the left that did not correspond to any ground truth event in either the left or unknown category. Using these definitions, we computed precision, recall, and the F1 score for the annotated hours (Figure S1C).

#### Curation of vocalizations

Detections were manually curated using a customized spectrogram browser adapted from the open-source MATLAB graphical user interface DeepSqueak (9). The browser was modified to display, edit, and export the associated detections for two aligned audio files. Curation involved correcting biotic sound boundary errors and removing abiotic false positives. Particular attention was given to correcting the boundaries of quiet but rapidly emitted vocalizations, which are especially challenging to segment automatically. PAIRId assignment detections were further reviewed to resolve “unassigned” vocalizations wherever possible. Four curated hours of PAIRId data with high vocal activity and different individual mice were designated as “ground truth” for evaluating assignment methods. Every hour from each dataset was curated, except for the long-term TeLC pre- and post-perturbation recordings, where only select hours were curated for the focal mouse.

#### Characterization of vocalizations

Vocalizations were characterized at both the individual note and temporal patterning levels. At the note level, several acoustic features were computed, including note duration, note amplitude, and pitch. Note duration was directly extracted from the detections. Note amplitude was determined as the peak amplitude within the 10 kHz to 120 kHz frequency range; when applicable, measurements from both high- and low-gain microphones were used. For the laryngeal phonation mechanism (heliox) experiments, the fundamental frequency was further quantified. For each note, a human annotator visually inspected the spectrogram and identified the lowest continuous trace and recorded the highest pitch of that trace. At the temporal patterning level, we first computed note-to-note temporal features. These features included the inter-event interval (the time between the end of the current event and the start of the next event), the log-transformed inter-event interval, the inter-start interval (the time between the start of the current event and the start of the next event), and the instantaneous rate (defined as the reciprocal of the inter-start interval). Next, to identify supra-note patterns of arbitrary length, we applied the following heuristics. First, a fixed-size 7-note sliding window was applied across all vocalizations. Within each window, two metrics were computed: the maximum log-transformed inter-event interval (representing the largest temporal gap) and the normalized root mean square error (NRMSE) from a linear regression of instantaneous rate versus note index. A window was deemed temporally patterned if it exhibited both a low maximum log inter-event interval (*≤* 100 ms), indicating temporal contiguity, and a low NRMSE (*≤* 0.15), reflective of stereotyped note sequencing. Consecutive windows meeting these criteria were merged into longer segments - either through direct overlap or, when non-overlapping, if the time gap between them did not exceed 0.5 seconds - to form supra-note patterns. Each resulting segment was then further characterized using RANSAC regression to robustly fit a linear model that accounted for outliers, thereby enabling the identification of long patterns with simple linear stereotypy. Once long patterns were delineated, we extracted key patterning features - specifically, the maximum rate (*r*_*max*), minimum rate (*r*_*min*), and the number of notes from the RANSAC fit. These features were subjected to principal component analysis (PCA) for dimensionality reduction to 2D, and a Gaussian mixture model (GMM) was then applied to cluster the segments into three distinct groups. Each note was subsequently assigned a category based on whether it belonged to a segment in one of the three patterning clusters or was unpatterned. Finally, by examining both note amplitude and spectrograms, we defined the four categories: songs, semi-songs, and patterned and unpatterned USVs. See Figure S2 for visuals accompanying these methods. To further characterize the temporal properties of songs and USVs, we also computed an alternative, complementary method for defining vocal bouts. Vocal bouts were defined as sequences of notes with inter-event intervals shorter than 100 ms, consistent with the definition of “group” in (10). Using this criterion, we identified song bouts and USV bouts and extracted their durations.

#### A mathematical model of the song rhythm

The song is composed of a series of progressively longer notes that evolve predictably over 6–10 seconds. We derived a simple mathematical model of the stereotyped temporal patterning of the song. We observed that the instantaneous note rate *r*(*i*) decreases linearly with the note index *i*, and postulated the linear model:

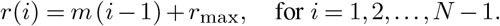

where

- *r*_max_ = *r*(1) is the maximum instantaneous rate (i.e. the start rate),
- *m <* 0 is the slope (indicating a decrease in rate as *i* increases),
- *N* is the number of notes.

Then, we also defined the minimum rate by extrapolating this linear relationship to the last note:

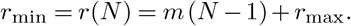

Thus, the three parameters *r*_*max, r*_*min*, and *m* fully characterize the temporal patterning of a song. From these three parameters, we next derived the total duration of the song *T*, a global property of the song. The total duration *T* (i.e., the time difference between the first and last note) is given by

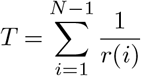

To obtain an analytic expression, we approximated this sum by the integral

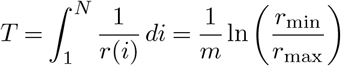

Since *m <* 0 and *r*_min_*/r*_max_ *<* 1, the logarithm is negative, and division by the negative *m* yields a positive duration *T*. Alternatively, one may write:

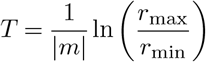

Thus, this model enabled us to analyze song variability (e.g., duration) in terms of these three interpretable parameters.

#### Video analysis

Using the overhead camera synchronized with audio in the PAIRId social assay enclosures, we quantified inter-animal distance for each dyad. We preprocessed videos of each enclosure by correcting the lens distortion using calibrated camera parameters. To estimate pose, we generated a SLEAP model using 950 manually labeled training frames and the multi-animal top-down pipeline with a single instance (11). The model estimated a skeleton with 6 points (nose, each ear, back of head, middle of spine, base of tail). The output nodes were then converted from pixel space in each video to common world coordinates. The middle-of-spine node was considered the centroid of each animal and used to calculate inter-animal distance. See Figure S1A for a visual overview.

#### Social interactions in the PAIRId setup

To determine if the two mice are socially engaged in the PAIRId setup, we first compared the number of vocalizations in the PAIRId setup to those in the “alone” condition. We defined “socially active” hours as those with note counts exceeding the threshold of the mean plus three times the standard deviation in alone hours. For these socially active hours, we further examined both the spatial and temporal organization of vocalizations. Spatially, we examined the locations of the mice in common world coordinates during vocalizations. Temporally, we computed the vocalization rate for each side using a rolling window (30-second window with a 15-second step size) and then calculated the Pearson correlation between the rates from the two sides. We also generated shuffled controls by mismatching the left and right rates from different hours for comparison.

#### Respiration analysis

To identify respiratory events, we analyzed the signal from the intranasal thermistor. The raw signal was first downsampled to 500Hz and then filtered using a 4th order Butterworth bandpass filter (0.5 - 50Hz). Inhalation onsets were detected as prominent peaks (high temperature), while offsets (marking the start of exhalation) were identified as corresponding troughs (low temperature). Putative respiratory cycles were defined by pairing each inhalation onset with the unique subsequent inhalation end occurring between consecutive onsets. Cycles with a duration shorter than 0.5 seconds were considered valid and retained for downstream analysis.

